# Acetylation of Axonal G3BP1 through ELP3 Accelerates Axon Regeneration

**DOI:** 10.1101/2025.10.28.685144

**Authors:** Irene Dalla Costa, Emily Michenfelder, Sophia Siciliano, Anika Tapita, Courtney N. Buchanan, Lauren S. Vaughn, Jinyoung Lee, Juan Oses-Prieto, Chunlong Ma, Elizabeth Thames, Nitzan Samra, Shifra Ben-Dor, Rebecca Haffner-Krausz, Mary McElveen, Terika P. Smith, Nabanita Nawar, Pimyupa Manaswiyoungkul, Patrick T. Gunning, Mike Fainzilber, Alma L. Burlingame, Haining Zhu, Eran Perlson, Pabitra K. Sahoo, Jeffery L. Twiss

## Abstract

Nerve injury triggers localized translation of axonal mRNAs to respond to injury and nerve regeneration. The core stress granule protein G3BP1 sequesters axonal mRNAs in granules before and after axotomy. G3BP1 granule disassembly can be regulated by post-translational modifications, including phosphorylation of S149 phosphorylation and acetylation of human K376 (mouse K374). Axonal G3BP1 undergoes phosphorylation after axotomy, but acetylation of G3BP1 in axons was unknown. Here we show that rodent G3BP1 undergoes K374 acetylation after axotomy is ELP3-dependent, which enhances axonal protein synthesis, accelerates nerve regeneration, and supports functional recovery. ELP3-depleted neurons exhibit reduced axon growth and increased axonal G3BP1 granules. The proximal axons degenerate rapidly despite maintaining soma connectivity, an effect prevented by expression of acetylmimetic G3BP1.Together, these findings identify G3BP1 acetylation via ELP3 as a critical regulator of both axonal regeneration and neuronal resilience, revealing a post-translational mechanism that links stress granule regulation to neuronal repair and protection.

## INTRODUCTION

Neurons in the peripheral nervous system (PNS) can spontaneously regenerate injured axons. This is due to a regeneration-permissive environment as well as a high intrinsic growth capacity that derives in part from a rapid activation of protein synthesis in injured axons. The locally synthesized proteins provide axon-to-soma retrograde signals and promote growth cone formation to initiate regeneration^1^. Stress granules (SG) are dynamic, membraneless cellular compartments composed of RNA binding proteins (RBPs) and translationally-inactive mRNAs that form by liquid-liquid phase separation (LLPS) during periods of cellular stress to store unneeded mRNAs^2^. The SG protein G3BP1 is used to store mRNAs in axons, and disassembly of axonal G3BP1 granules promotes axon regeneration in both the PNS and central nervous system (CNS)^3–5^. G3BP1 phosphorylated on Serine 149 (G3BP1^PS149^) has an increased threshold for liquid-liquid phase separation (LLPS)^6,7^. In axons, casein kinase 2α (CK2α) phosphorylates G3BP1 S149 shortly after axotomy, an event that decreases the abundance of G3BP1 granules in axons, increases axonal protein synthesis by releasing mRNAs from storage, and promotes axon regeneration^8^. Phosphorylation of G3BP1 requires a sequential activation of protein synthesis in axons where *mTor* mRNA is translated first and its protein product then activates local synthesis of CK2α^4,9^. G3BP1 is known to undergo other post-translational modifications, including ubiquitination at K63 and acetylation at human K376 (mouse and rat K374)^10,11^. While K63 ubiquitination promotes disassembly of G3BP1 granules^11^, K376 acetylation of human G3BP1 (hG3BP1) decreases the protein’s RNA binding activity^10^ and RNA binding decreases G3BP1’s threshold for LLPS^6,7^. Human G3BP1 K376 and mouse G3BP1 K374 are within G3BP1’s RRM RNA binding domain, so acetylation promotes disassembly of SGs^10^.

Though PNS nerves spontaneously regenerate, they grow very slowly at 1-3 mm/d^12–15^ and regeneration of distances of more than 5-6 cm are typically unsuccessful. This failed regeneration is in large part due to a decline in growth promotion/permissiveness distal to the injury site^16^. Thus, there is a clear need for axon growth-accelerating therapies for PNS injury. Considering our previous studies showing that G3BP1 granules slow axon regeneration^3–5^, we asked if acetylation of axonal G3BP1 contributes to PNS axon regeneration. We show that rodent G3BP1 can be acetylated at K374 (G3BP1^AcK374^) in sciatic nerve axons after crush injury. Expression of acetylmimetic human G3BP1 (hG3BP1^K376Q^) increases axon growth in cultured rodent dorsal root ganglion (DRG) neurons and regeneration in injured sciatic nerve in vivo, an effect accompanied by increased protein synthesis in axons but not in the soma.

Previous studies reported Histone Deacetylase 6 (HDAC6) and CREB Binding Protein (CREBBP or CBP) as hG3BP1 deacetylase and lysine acetyl transferase (KAT), respectively^10^. We confirmed that indeed mouse HDAC6 deacetylates axonal G3BP1, but found that the axonal G3BP1 KAT is actually Elongator complex protein 3 (ELP3). Depleting ELP3 from neurons increases axonal, but not cell body, G3BP1^AcK374^ and increases the number of axonal G3BP1 granules to slows axon growth in cultured rodent DRG neurons. Surprisingly, proximal axons of ELP3-depleted neurons degenerate after severing in culture and in vivo axotomy, which is mitigated by acetylmimetic G3BP1.

Taken together, these findings establish G3BP1 acetylation as a key post-translational mechanism that simultaneously promotes axonal regeneration and protects axons from stress-induced degeneration. ELP3 levels are reduced in post-mortem spinal cord from amyotrophic lateral sclerosis (ALS) and brain from Alzheimer’s disease patients^17,18^, suggesting that impaired G3BP1 acetylation may contribute to pathological stress granule accumulation and neuronal vulnerability in neurodegenerative disorders. Modulating ELP3 activity or mimicking G3BP1 acetylation may therefore provide a therapeutic strategy to enhance regeneration while preserving axonal integrity.

## RESULTS

### Axonal G3BP1 is rapidly acetylated following axotomy

G3BP1 forms SG-like structures in naïve axons^19^. Shortly after injury, these axonal G3BP1 granules increase in number and size, and subsequently decrease without completely disappearing during regeneration^3,5^. Axons show a commensurate increase in G3BP1^PS149^ concomitantly with the decrease in axonal G3BP1 granules^8^. Gal et al. (2019) reported that G3BP1 can be acetylated in U2OS cells but the possibility that this occurs in axons had not been tested^10^. Thus, we asked whether G3BP1 undergoes K374 acetylation in rodent peripheral nerve using an anti-human G3BP1^AcK376^ antibody for immunofluorescent detection^10^. Anti-G3BP1^AcK376^ signals were barely detectable in axons before injury, with a sharp rise at 3 h that decreased to near naive levels by 16 h after crush lesion (Figure 1a,b). Interestingly, the rise in axonal G3BP1^AcK374^ roughly paralleled the increase in axonal G3BP1^PS149^ (Figure 1c,d). Co-labeling with anti-G3BP1 and anti-acetyl-lysine (Ac-Lys) antibodies, followed by stimulated energy depletion (STED) confocal imaging, similarly showed an increase in colocalizing signals at 3 h after nerve crush indicative of G3BP1 acetylation (Figure 1e,f). M1 and M2 coefficients indicated significantly more G3BP1 colocalizing with Ac-Lys in injured axons, but Ac-Lys colocalizing with G3BP1 was not signficant consistent with more axonal acetylated proteins beyond G3BP1 in the injured axons (Figure 1g,h). Taken together, these data indicate that PNS nerve injury stimulates G3BP1 acetylation in axons.

**Figure 1:**
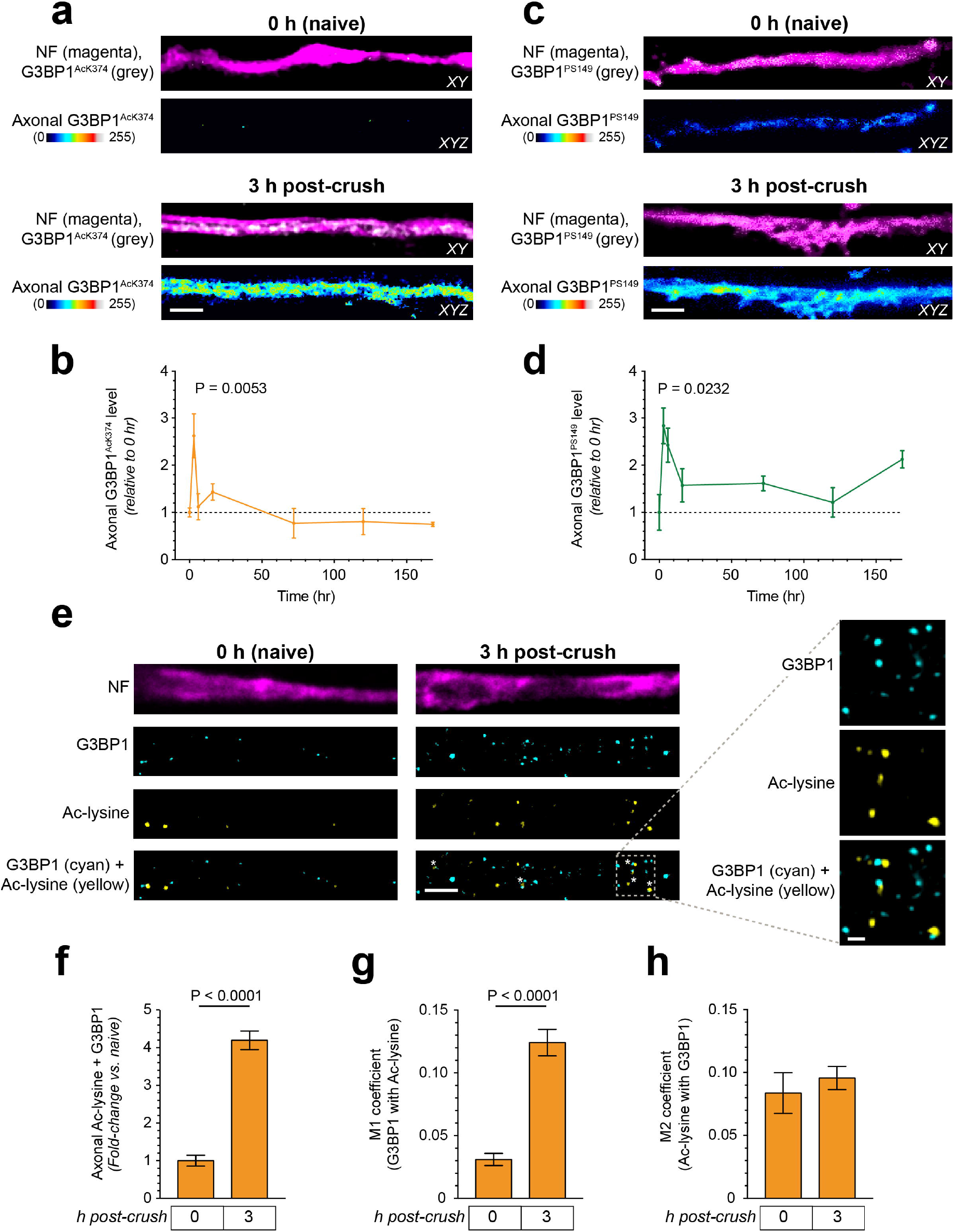
Axotomy increases acetylation of axonal G3BP1. **a-d)**, Representative confocal images of sciatic nerve axons for neurofilament (NF) and G3BP1^AcK374^ (**a**) or G3BP1^PS149^ (**c**) at 0 and 3 h post-crush injury taken at 0.5-1.5 mm proximal to the injury site are shown. Upper panel of images pairs shows XY plane through axon and lower panel shows axonal G3BP1^AcK374^ as maximal XYZ projection generated from G3BP1^AcK374^ pixels that overlapped with NF signal across individual Z planes. Quantitation of axonal G3BP1^AcK376^ (**b**) or G3BP1^PS149^ (**d**) levels relative are shown to naive as mean ± standard error of the mean (SEM) [Scale bar = 5 µm]. **e-h)**, Representative STED images for G3BP1 and Acetyl-lysine (Ac-Lys) antibodies in axons proximal to the injury site as in panel a (**e**). Inset panels show high magnification for indicated region [Scale bars = 5 µm main panels, 1 µm insets]. Quantification of G3BP1 and Ac-Lys signal colocalization is shown in **f** based mean fold-change ± SEM with mean ± SEM for M1 (**g**) and M2 (**h**) coefficients shown (N = 3-7 animals per condition for b, d and f-h; P values are determined by one-Way ANOVA with Tukey’s post-hoc test for b and d and student’s T-test for f-h).

### Acetylmimetic G3BP1 increases axon growth

G3BP1 S149 phosphorylation causes axonal G3BP1 granule disassembly, decreases RNA binding by G3BP1, and promotes axon growth^4,5^. To determine if G3BP1 acetylation affects axon growth, we introduced mCherry (mCh) fused to wild type, acetylmimetic and non-acetylatable hG3BP1 (hereafter hG3BP1 and mutant variants) into dissociated DRG neuron cultures and analyzed axon growth. For this, we generated K376 to glutamine and arginine mutations in hG3BP1-mCh (acetyl-mimetic, hG3BP1^K376Q^-mCh, and nonacetylatable, hG3BP1^K376R^-mCh, respectively); wild type (WT) hG3BP1-mCh and mCh were used as controls (Figure 2a). hG3BP1^K376Q^-mCh expressing neurons showed significantly longer axons than the hG3BP1^K376R^-mCh, hG3BP1-mCh, and mCh expressing neurons (Figure 2b,c; Supplementary Fig. 1a). Axon branching did not change significantly for any of hG3BP1 compared to mCh transfected neurons; however, there were modest changes in branching comparing between the mutant and WT hG3BP1 transfected DRGs (Supplementary Fig. 1b,c). Expression of the non-acetylatable hG3BP1^K376R^-mCh did not decrease axon growth (Figure 2b,c) unless endogenous G3BP1 was depleted (Figure 2d-f and Supplementary Fig. 1d,e and 2a-c). Note that the siRNAs used to deplete endogenous G3BP1 target the 3’ untranslated region (UTR) of *G3bp1* mRNA, target sequences not included in the hG3BP1-mCh expression constructs.

**Figure 2:**
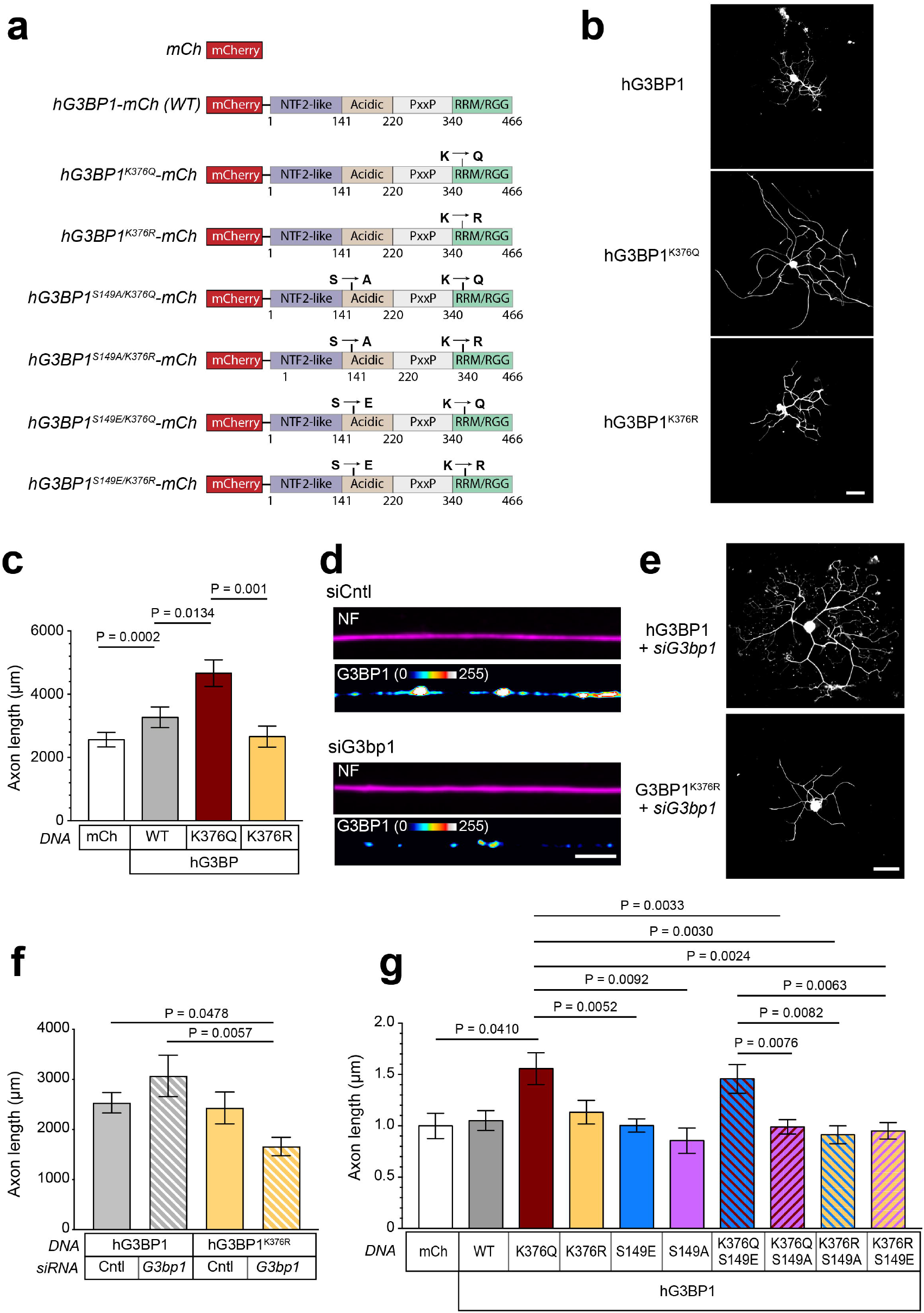
Acetylmimetic G3BP1 increases axon growth. **a)**, Summary of wild type (WT) and mutant human G3BP1-mCh (hG3BP1) constructs used in b-g with defined G3BP1 domains designated. **b-c**), Representative images of NF signals in dissociated DRG neuron expressing WT and K376 mutants (**b**). Quantitation of mean axon length per neuron ± SEM is shown for DRG cultures from b after 65 h in culture (c); see Supplementary Figure 1a-c for axon length histogram, mean longest axon, and axon branching [Scale bar = 200 µm]. **d)**, Representative images of axons for DRG neurons transfected with non-targeting or G3bp1 mRNA targeting siRNAs; see Supplementary Figure 1d,e for (siCntl and siG3bp1, respectively) [Scale bar = 10 µm]. **e-f)**, Representative images for NF in DRG neurons co-transfected with siG3bp1 + WT or mutant hG3BP1 shown in e. Mean axon lengths per neuron ± SEM for DRGs co-transfected as in e at 65 h in culture are shown in **f**; see Supplementary Figure 2a-c axon length histogram, mean longest axon, and axon branching [Scale bar = 200 µm]. **g)**, Mean axon length ± SEM for expressing mCherry (mCh), WT hG3BP1 or indicated single and double hG3BP1 mutants; see Supplementary Figure 2d,e for longest axon and axon branching data (N ≥ 70 neurons over 3 independent cultures in c, ≥ 57 in f and g ≥ 102; P values by two-way ANOVA with Tukey’s post-hoc test in c, f and g).

Since G3BP1 S149 phosphorylation also causes G3BP1 granule disassembly, we asked if dual phosphomimetic and acetylmimetic hG3BP1 might provide a synergistic or additive effect over single mutants. Thus, we generated phosphomimetic + acetylmimetic hG3BP1^S149E/K376Q^-mCh, non-phosphorylatable + acetylmimetic hG3BP1^S149A/K376Q^-mCh, phosphomimetic + non-acetylatable hG3BP1^S149E/K376R^-mCh, and non-phosphorylatable + non-acetylatable hG3BP1^S149A/K376R^-mCh expressing constructs (Figure 2a). Axon length in the hG3BP1^S149E/K376Q^-mCh expressing neurons was no greater than the acetylmimetic hG3BP1^K376Q^ (Figure 2g). The growth-promoting effect of acetylmimetic hG3BP1 was lost when the non-phosphorylatable S149A mutation was included; likewise, the phosphomimetic S149E mutation did not increase axon growth in the context of the non-acetylatable K376R mutation (Figure 2g; Supplementary Fig. 2d,e). These data indicate that G3BP1 acetylation more strongly promotes axon growth than G3BP1 phosphorylation. Since the S149A mutation attenuated the effects of the acetylmimetic mutation, we cannot exclude the possibility that initial phosphorylation impacts functionality of the acetylated G3BP1. Though we saw overall little axon growth effects by the single S149 phosphomimetic and non-phosphorylatable hG3BP1 mutants (Figure 2g; Supplementary Fig 3d,e), preventing G3BP1 S149 phosphorylation by CK2α depletion decreased axon regeneration^8^, indicating both acetylation and phosphorylation of G3BP1 are needed for axon regeneration.

### Acetylmimetic G3BP1 increases axonal protein synthesis

Expression of G3BP1’s intrinsically disordered region 1 (IDR1; amino acids 141-220) and treatment with a cell permeable peptide containing rodent G3BP1 amino acids 190-208 disassemble G3BP1 granules to increase axonal protein synthesis^3,5^. Further, Gal et al. (2019) showed that G3BP1 acetylation disassembled G3BP1 granules in U2OS cells^10^. Thus, we asked if acetylmimetic G3BP1 affects axonal protein synthesis using the puromycin analog O-propargyl-puromycin (OPP) to label nascent peptides by Click-It chemistry^20^ in the hG3BP1^K376Q^-mCh, hG3BP1^K376R^-mCh, hG3BP1-mCh, and mCh transfected DRG neuron cultures (Figure 3a). hG3BP1^K376Q^-mCh expressing neurons showed significantly higher OPP incorporation in axons than hG3BP1^K376R^-mCh, hG3BP1-mCh and mCh expressing neurons (Figure 3b,c). However, this effect was limited to axons, as there were no significant differences in OPP incorporation in the soma of hG3BP1 WT and mutants and mCh expressing neurons (Figure 3d,e). Protein synthesis inhibition with anisomycin confirmed that the OPP signals are from newly synthesized proteins (Figure 3b-e). Thus, faster axon growth in DRG neurons expressing acetylmimetic G3BP1 is accompanied by higher protein synthesis in the growing axons.

**Figure 3:**
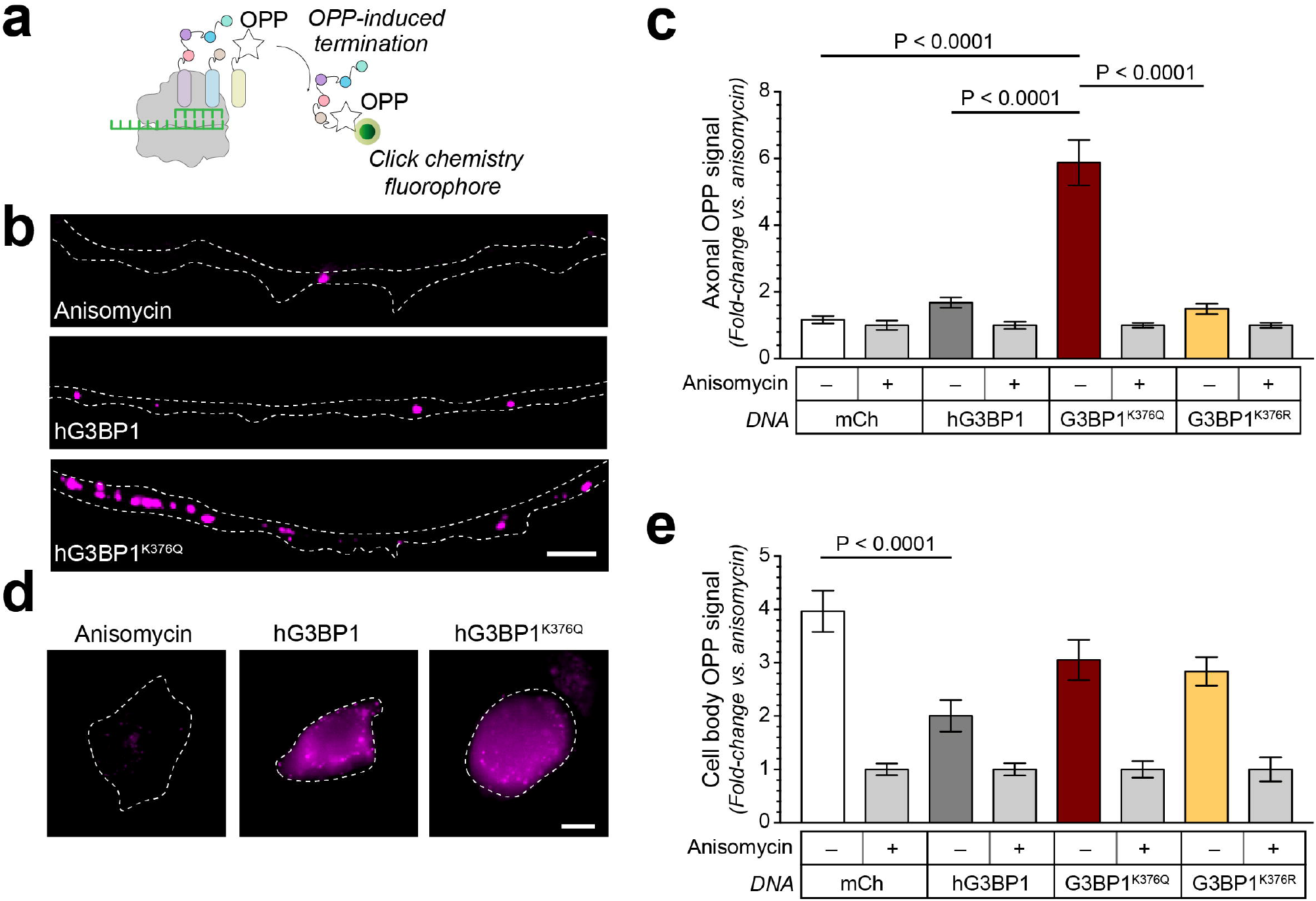
Acetylmimetic G3BP1 increases axonal protein synthesis. **a)**, Schematic representation of OPP (O-propargyl puromycin) labelling for detection of nascent protein synthesis. **b)**, Representative images of OPP signals in axons of DRG cultures expressing WT vs. mutant hG3BP1 shown in **b** at 65 h in culture. Anisomycin treated transfected cultures was used to verify OPP signals represent protein synthesis. **c** shows quantification of the axonal OPP signals for indicated constructs ± anisomycin [Scale bar = 5 µm]. **d-e)**, Representative images for OPP signals in soma of DRGs as in **b** [Scale bar = 5 µm]. **e** shows mean ± SEM of the soma OPP signals for indicated constructs ± anisomycin (N ≥ 104 axons in c and 16 neurons in **e** across 3 independent cultures; P values by two-way ANOVA with Sidak correction).

### Acetylmimetic G3BP1 accelerates peripheral nerve regeneration

Given the axon growth promotion and increased axonal protein synthesis seen with acetylmimetic G3BP1 expression, we next asked if G3BP1 acetylation might increase axon regeneration *in vivo* after peripheral nerve injury. For this, we generated adeno-associated serotype 9 viruses (AAV) expressing hG3BP1^K376Q^-mCh, hG3BP1^K376R^-mCh, hG3BP1-mCh or mCh. AAVs were injected into the proximal sciatic nerve of mice and mid-thigh nerve crush was performed 14 d later. We have previously shown that nerve-injected AAVs are retrogradely transported for expression of encoded cDNAs in sensory and motor neurons^5,21^. Morphological analyses of sciatic nerves at 7 d post-injury showed higher regeneration indices for the AAV-hG3BP1^K376Q^-mCh transduced mice compared to those transduced with AAV-hG3BP1^K376R^-mCh, -hG3BP1-mCh or -mCh (Figure 4a,b).

**Figure 4:**
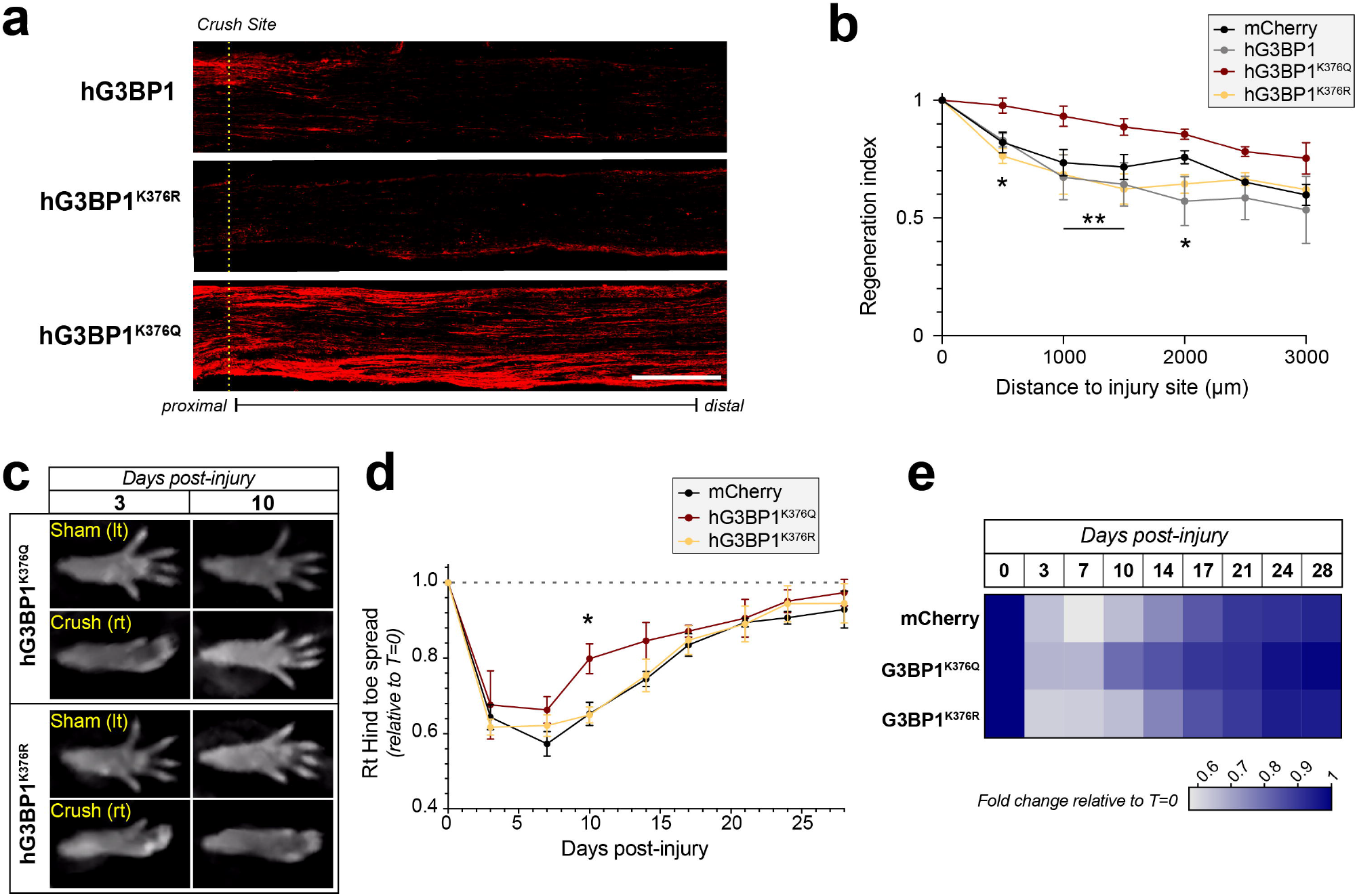
G3BP1 acetylmimetic increases sciatic nerve regeneration. **a)**, Representative widefield images for SCG10 immunofluorescence as a marker for regenerating axons in sciatic nerves of mice transduced with AAV-hG3BP1, -hG3BP1^K376R^, or -hG3BP1^K376Q^ at 7 d following nerve crush (21 d post-AAV transduction). Dashed line indicates crush site [Scale bar = 500 µm]. **b)**, Quantification of the regeneration indices from panel a expressed as mean fraction of SCG10 axon profiles at indicated sites relative to immediately proximal to the crush site ± SEM. c), Representative images of hind toe spread at 3 and 10 d after right sciatic nerve crush injury in mice transduced with AAV-mCh or AAV-hG3BP1 mutants. ‘Crush’ indicates hind paw ipsilateral to crush injury (rt) and ‘sham’ is hind paw contralateral to crush injury (lt). **d-e)**, Ratio for crush to sham (rt/lt) hind paw toe spread ± SEM is shown for indicated times post-crush injury for AAV-mCh, -hG3BP1^K376Q^, and -hG3BP1^K376R^ transduced mice as in a (**d**); see Supplementary Figure 3 for additional functional outcomes and Supplementary Video 1 for representative live imaging sequence. Heat map representation of toe spread ratios is shown in **f** (N = 4-6 mice in b and 5-6 in d and e; P values are determined by two-way ANOVA with Sidak correction in b, d and e; * p ≤ 0.05 and ** ≤ 0.01).

To assess the possibility of increased functional recovery from nerve injury with hG3BP1^K376Q^-mCh expression, we used non-invasive near-infrared light imaging to assess recovery of hind limb usage^22^. We bilaterally transduced with AAV by sciatic nerve injection to control for any nerve injuries inflicted by the intranerve injections. At 10 d after a right-sided unilateral nerve crush, significantly increased recovery of toe spread was seen in AAV-hG3BP1^K376Q^-mCh vs. -mCh and -hG3BP1^K376R^ transduced mice (Figure 4c-e; Supplementary Video 1). There was a trend for increased toe spread out to 17 d post-injury in the AAV-hG3BP1^K376Q^-mCh transduced mice compared to the AAV transductions, but this did not reach statistical significance (Figure 4d,e). hG3BP1^K376Q^ expression also increased the ratio of right hind paw to front paw usage compared to mCh and hG3BP1^K376R^ expression (Supplementary Fig. 3a,b). Right to left hind paw usage and standing ratios were also higher in the hG3BP1^K376Q^ vs. hG3BP1^K376R^ expressing mice (Supplementary Fig. 3c-f), but there was no difference in distance travelled by the mice between the different conditions (Supplementary Fig. 3g,h). Together, these data suggest that G3BP1 acetylation accelerates both morphological and functional PNS nerve regeneration.

### HDAC6 deacetylates and ELP3 acetylates G3BP1

Given the growth promoting effects of G3BP1 acetylation and the rapid appearance of this modification after peripheral nerve injury, we asked how this post-translational modification is modulated. HDAC6 was reported to be a G3BP1 deacetylase^10^, and HDAC6 localizes to peripheral axons where it has growth-attenuating effects^23,24^. To determine if HDAC6 deacetylates G3BP1 in axons, we treated rat DRGs cultures with the HDAC6 inhibitor Tubastatin A (TubA). DRGs neurons treated with TubA displayed an increased G3BP1^AcK374^ (Figure 5a), suggesting that HDAC6 can deacetylate neuronal G3BP1. Consistent with this increase in G3BP1^AcK374^ with HDAC6 inhibition, TubA also increased axon growth in the DRGs and this effect was lost when endogenous G3BP1 was depleted using siRNA (Figure 5b,c and Supplementary Fig. 4a-c). Similarly, a second HDAC6 inhibitor increased axon growth in DRG cultures (Supplementary Fig. 5a-d).

**Figure 5:**
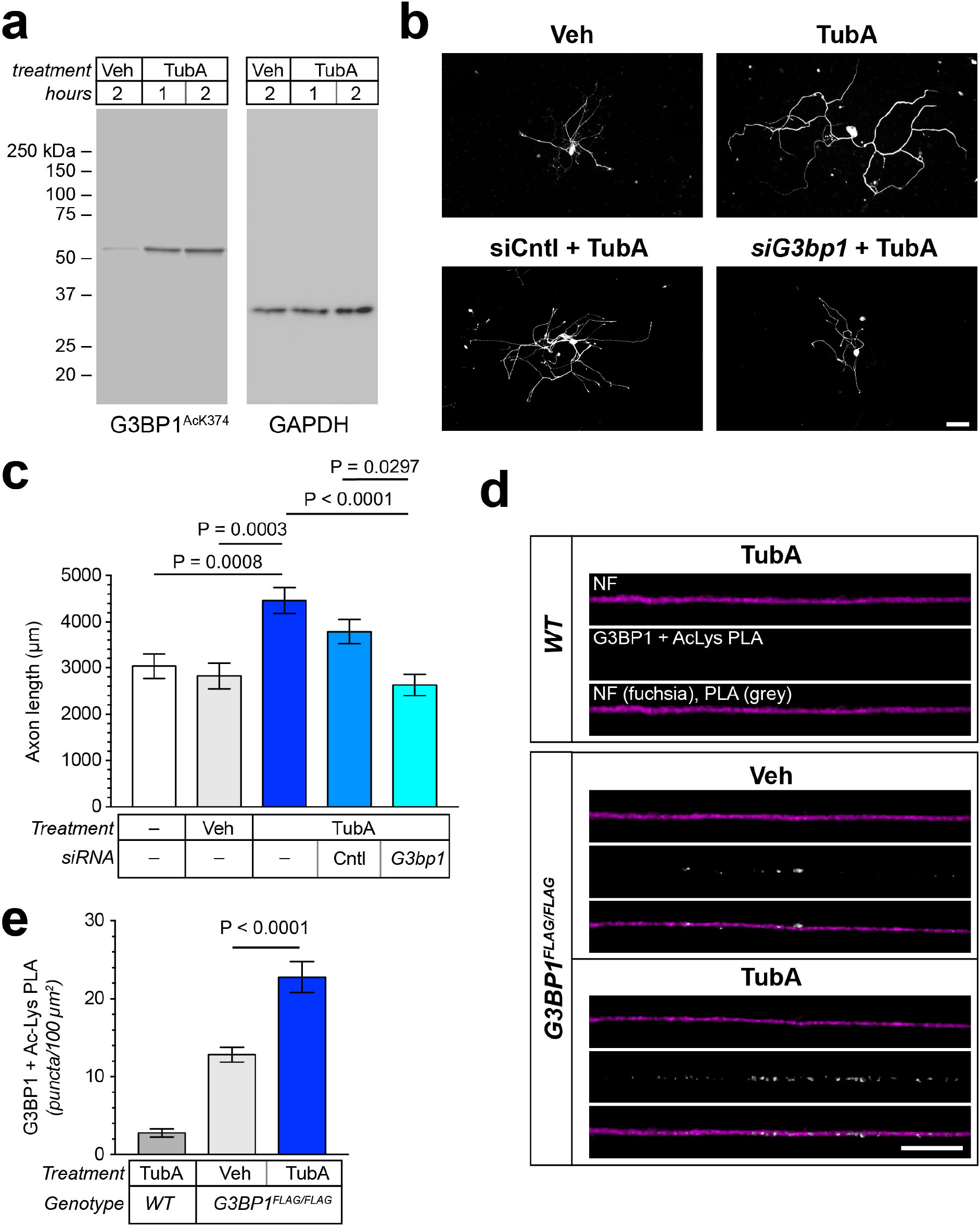
HDAC6 inhibition increases axonal G3BP1^AcK376^ and axon growth. **a)**, Immunoblot for anti-hG3BP1^Ac376^ using lysates from cultures of adult rat DRGs treated with 10 µM TubA for 1-2 h vs. vehicle control (DMSO). GAPDH shows equivalent loading between lanes. **b-c)**, Representative NF immunostaining of DRG neurons treated with 10 µM TubA for 2 h ± siG3bp1 or siCntl shown in **b**. Axon length per neuron shown as mean ± SEM in **c**. See Supplementary Figure 5 for growth data for DRGs treated with HDAC inhibitor NN-769 and see Supplementary Figure 4a-c for axon length histogram, mean longest axon, and axon branching [Scale bar = 200 µm]. **d-e)**, Representative images for anti-Ac-Lys + -FLAG (for G3BP1^FLAG^) PLA signals in axons of DRGs cultured from WT vs. G3BP1^FLAG/FLAG^ mice DRG neurons treated with 10 µM TubA for 2 h vs. vehicle control (**d**) [Scale bar = 200 µm]. Quantification of axonal PLA puncta across 100 µm^2^ axon area from DRG cultures treated as in d shown in **e**; see Supplementary Figure 4d,e for FLAG mice validation (N ≥ 116 axons in c and ≥ 160 in e across 3 independent cultures; P values by two-way ANOVA with Tukey’s post-hoc test).

To more directly assess whether blocking HDAC6 increases G3BP1^AcK374^ we used a proximity ligation assay (PLA) with anti-Ac-Lys and anti-FLAG antibodies. We specifically designed mice with a FLAG-tagged G3BP1 allele (G3BP1^FLAG/FLAG^) to gain higher affinity detection of G3BP1 than achieved with available anti-G3BP1 antibodies. Monoclonal anti-FLAG antibodies proved highly specific by immunofluorescence, western blots, and immunoprecipitations as a proxy for G3BP1 in these mice (validated in Supplementary Fig. 4d,e). PLA puncta were clearly visible for anti-Ac-Lys + anti-FLAG along axons of DRGs cultured from G3BP1^FLAG/FLAG^ but not WT mice, and the axonal PLA signal for acetylated G3BP1^FLAG/FLAG^ in DRG cultures was significantly higher after TubA treatment (Figure 5d,e). Thus, HDAC6 inhibition promotes axon growth and increases axonal G3BP1 acetylation.

CREBBP is known to acetylate histones in the nucleus as well as some cytoplasmic proteins^25^ and it was reported to acetylate G3BP1 in U2OS cells^10^. We searched a rat sciatic nerve axoplasm proteomics dataset for KATs localizing into PNS axons; tandem mass tagging was used in that proteomics experiment to compare naïve and regenerating axons. We found that the KATs α-tubulin acetyl-transferase 1 (ATAT1)^26^, CREBBP^25^, and ELP3^27,28^ are present in the axoplasm based on the detection by mass spectrometry of ions with fragmentation spectra identifying specific peptides of these three acetyl transferases, in the tryptic digests of these samples (Supplementary Fig. 6). We individually depleted *Atat1, Crebbp*, and *Elp3* mRNAs from DRG cultures using siRNAs and asked if depletion affects axonal G3BP1 granules and G3BP1^AcK374^ levels. Reverse transcriptase-coupled droplet digital PCR (RTddPCR) confirmed depletion of the endogenous *Atat1, Crebbp* and *Elp3* mRNAs compared to siCntl-transfected cultures (Figure 6a; Supplementary Fig. 7a-c). The Elp3-depleted neurons, but not the *Atat1*- or *Crebbp*-depleted neurons, showed a clear decrease in axonal G3BP1^AcK374^ levels compared to siCntl-transfected DRGs (Figure 6b,c). Interestingly, there was no change in G3BP1^AcK374^ levels in the cell bodies of these neurons (Supplementary Fig. 7d). DRG neurons from G3BP1^FLAG/FLAG^ mice similarly showed a decrease in acetylated axonal G3BP1 by PLA assay (Supplementary Fig. 7e,f). Consistent with the effects of *Elp3* depletion above, *siElp3* transfection decreased axon growth compared to siAtat1, siCrebbp, and siCntl transfected neurons (Figure 6d,e; Supplementary Fig. 7g). *Elp3* depletion also increased axonal G3BP1 granules compared (Figure 6f-g). These data suggest that ELP3 serves as a G3BP1 KAT in DRG axons.

**Figure 6:**
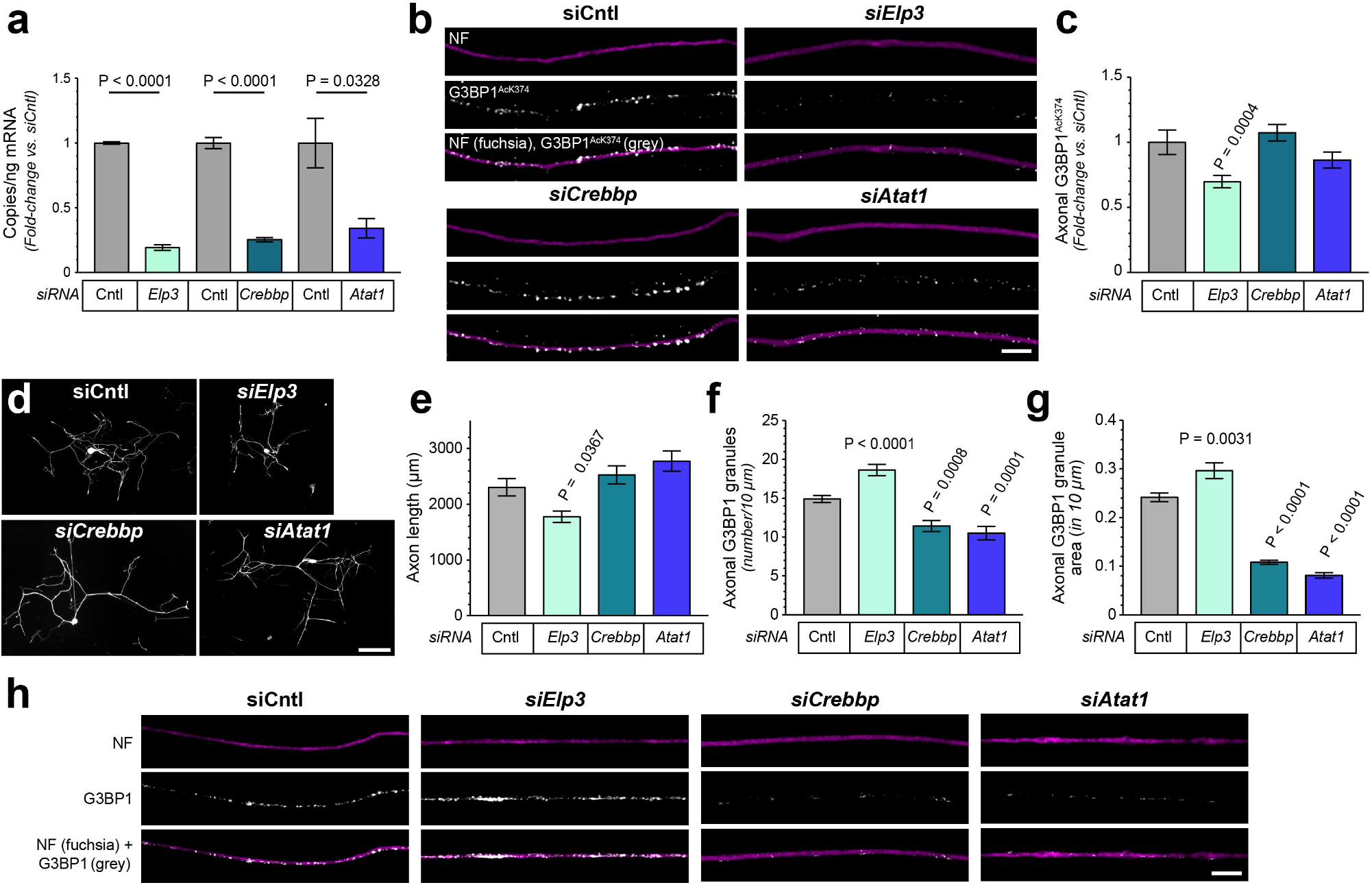
Axonal G3BP1 is acetylated by ELP3. **a)**, RTddPCR for Elp3, *Atat1* and *Crebbp* mRNAs (normalized for *Hmgb1* values) in mouse dissociated DRG cultures transfected with indicated siRNAs shown as mean fold-change in copies/ng relative to control ± SEM. See Supplementary Figure 7a-c for axonal and soma ELP3 protein immunofluorescence in siCntl vs. siElp3 transfected DRG cultures. **b-c)**, Representative images for NF and G3BP1^AcK374^ in DRG neurons treated with siCntl, *siElp3, siCrebbp*, or *siAtat1* shown in **b**. Quantitation of axonal G3BP1^Ac374^ immunofluorescence shown in c as mean fold-change relative to siCntl ± SEM. See Supplementary Figure 7d for soma G3BP1^AcK374^ levels with these siRNAs; see Supplementary Figure 7e,f for PLA data [Scale bar = 5 µm]. **d-e)**, Representative images of NF immunostaining for dissociated DRG neurons transfected with indicated siRNA shown in **d**. Mean axon length/neuron for siRNA transfected cultures is shown in e; see Supplementary Figure 7g for longest axon and axon branching per neuron [Scale bar = 200 µm]. **f-g)**, Quantification of axonal G3BP1 granule number and density in siRNA transfected DRG cultures is shown as mean ± SEM (**f, h**). Representative NF + G3BP1 immunofluorescent signals are shown for axons of siRNA transfected DRG cultures shown in **g** (N = 3 cultures for a and ≥ 44 axons in c ≥ 36 axons in e, and ≥ 130 in g-h across 3 independent cultures; P values by one-way ANOVA with Tukey’s multiple post-hoc) [Scale bar = 5 µm].

### ELP3 depletion promotes degeneration of injured axons

We next asked if ELP3 affects axon regeneration. For this, we injected sciatic nerves of adult mice with AAV-shElp3-BFP or AAV-shCntl-EGFP respectively in the left and right sciatic nerve and then performed a bilateral sciatic nerve crush 14 d later at mid-thigh level about 0.5 cm distal from the injection site. Sciatic nerves were evaluated for regeneration at 5 d post crush using immunolabeling for SCG10. Mice transduced with AAV-shElp3-BFP displayed significantly lower regeneration indices than those transduced with AAV-shCntl-EGFP (Figure 7a; Supplementary Fig. 8a).

**Figure 7:**
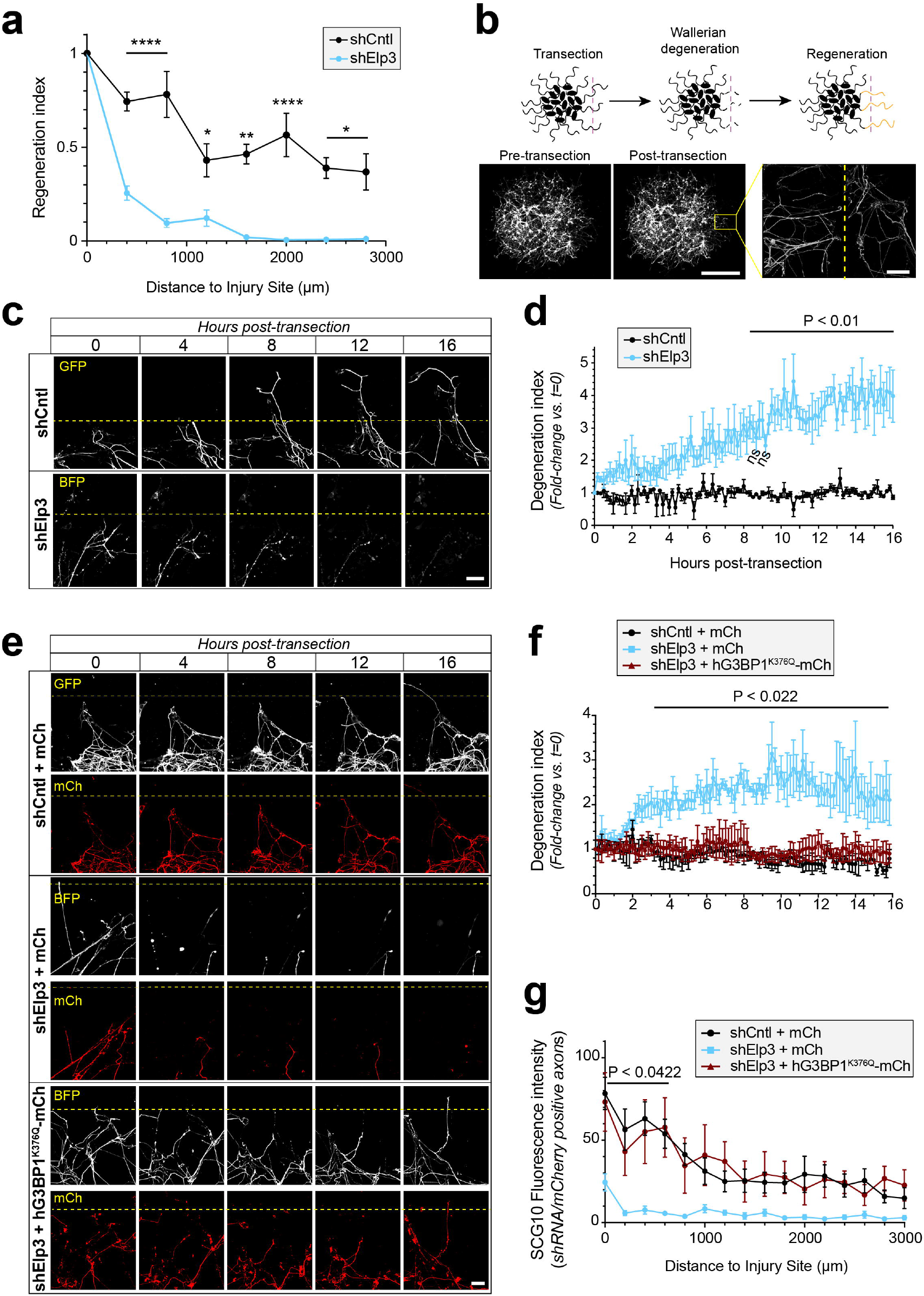
ELP3 depletion decreases sciatic nerve regeneration and leads to axonal degeneration. **a)**, Regeneration index quantitation based on SCG10 signal of sciatic nerve neurons transduced with AAV-shCntl-GFP vs -shElp3-BFP is shown as mean axon profiles ± SEM relative to immediately proximal to the crush site (see Supplementary Figure 8a for representative images). **b)**, Representative time course for regeneration in AAV-shCntl-GFP vs. -shElp3-BFP transduced mouse DRG spot cultures. Axons were transected after 7 d in culture and visualized immediately following transection by GFP and BFP signals. Dashed lines indicate transection site (above line is distal and below is proximal axon relative to transection). Note that ELP3-depleted proximal axons degenerate after transection; see Supplementary Video 2 for representative live imaging sequence and Supplementary Figure 8c,d for Degenotag analysis [Scale bar = 50 µm]. **c)**, Schematic for experimental design for the axonal regeneration from DRG ‘spot culture’ by manual transection of axon shafts (above) with representative low magnification images of spot cultures before and after transection (below). Inset panel shows high magnification of boxed region with dashed line indicating transected site [Scale bar = 200 µm in main panel and 10 µm for inset]. **d)**, Quantification of axon degeneration based on fragmentation over time is shown as mean degeneration index ± SEM for each time point. **e-f)**, Representative time course for regeneration in co-transduced with AAV-shCntl-GFP and -mCherry vs. AAV-shElp3-BFP and -mCherry vs AAV-shElp3-BFP and -mCherry-G3BP1^K376Q^ is shown for BFP/GFP and mCh signals in mouse DRG spot cultures as in c (**e**). Note that hG3BP1^K376Q^-mCh expression prevented degeneration of the ELP3-depleted proximal axons; see Supplementary Video 3 for representative live imaging sequence [Scale bar = 50 µm]. Degeneration indices (**f**) show hG3BP1^K376Q^ expression significantly prevents proximal axon degeneration seen after transection of ELP3-depleted DRG neurons **g)**, Quantitation of SCG10 fluorescence intensity of sciatic nerve neurons co-transduced with AAV-shCntl-GFP and -mCh vs. AAV-shElp3-BFP and -mCherry vs AAV-shElp3-BFP and -G3BP1^K376Q^-mCh is shown as mean fluorescence intensity ± SEM for each 200 µm bin from the crush site (Representative images in Supplementary Figure 8b; N = 5-6 mice for a, c and d, 4 for g and 3 for e,f; P values are determined by two-way ANOVA followed by Sidak correction in a, d, f and g).

The regeneration indices of the shElp3 expressing mice showed a sharp decline at ~400 µm distal to the crush site and very few axons were seen proximal to the injury site on close inspection of the BFP signals (Figure 7a, Supplementary Fig. 8a). To better visualize regenerating axons in real time when ELP3 is depleted, we turned to an *in vitro* axotomy model. For this, dissociated DRGs are transduced with AAVs and then cultured in a small ‘spot’ that allows for radiating axonal growth; axotomy was performed by manually transecting axons after 7 d in vitro^29^ (Figure 7b). As expected, both AAV-shElp3 and -shCntl transduced cultures showed degeneration of axons distal to the cut (Figure 7c; Supplementary Video 2). For AAV-shCntl transduced cultures, the proximal severed axons started to regenerate and cross the injury site over the 16 h time-lapse sequence (Figure 7c; Supplementary Video 2, upper panel). Although ELP3 depleted neurons are clearly capable of initially extending axons in culture, the proximal axons of the ELP3-depleted cultures (i.e., soma connected) degenerated after axotomy (Figure 7c,d; Supplementary Video 2, lower panel). To determine if G3BP1 contributes to the axon degeneration after injury in the ELP3-depleted neurons, we co-transfected AAV-siElp3-BFP transduced DRG spot cultures with AAV-hG3BP1^K376Q^-mCh (Figure 7e,f). Acetylmimetic G3BP1 expression attenuated proximal axon degeneration in the ELP3-depleted neurons (Figure 7e,f; Supplementary Video 3). This indicates that the axon degeneration seen with ELP3 depletion is cell autonomous, similar to Wallerian degeneration.

We finally asked if acetylmimetic G3BP1 might also rescue axons after *in vivo* axotomy in ELP3-depleted mice by co-transduction with AAV-shElp3 + AAV-mCh or AAV-hG3BP1^K376Q^-mCh. Given the rapid decline in regeneration indices seen in shElp3 transduced neurons in Figure 7a and Supplementary Fig. 8a, we reasoned that evaluating anti-SCG10 signal intensity in shRNA expressing (AAV-shCntl-GFP vs. AAV-shElp3-BFP) axons that also showed hG3BP1^K376Q^-mCh or mCh expression would allow us to capture axons of the dually transduced neurons. Acetylmimetic G3BP1 expression rescued the apparent axon degeneration seen with ELP3 depletion but also promoted axon growth across the injury site (Figure 7g and Supplementary Fig. 8b). To determine if sciatic nerve axons were indeed degenerating *in vivo* after the axotomy, we used *DegenoTag* antibody that specifically recognizes an epitope in cleaved low molecular neurofilament (NF⍰L) exposed in degenerating axons^30^. Sciatic nerves of AAV-shELP3 + AAV-mCh transduced mice showed significantly higher DegenoTag signals proximal to the crush site compared to AAV-shCntl + AAV-mCh and AAV-shELP3 + AAV-hG3BP1^K376Q^-mCh transduced mice, but equivalent signals distal to the crush site (Supplementary Fig. 8c,d). Taken together, these findings indicate that depletion of ELP3 places axons at risk for degeneration when exposed to a mechanical trauma despite still being connected to the soma. These data suggest that ELP3-dependent acetylation of axonal G3BP1 maintains the integrity of injured axons in addition to promoting their regeneration.

## DISCUSSION

Injured PNS axons regenerate slowly at approximately 1-3 mm per day^12–15^ and regeneration distances over 5-6 cm frequently result in incomplete functional recovery^16^. Accelerating nerve regeneration is a critical goal for restoring function after traumatic PNS injury^31^. Here, we show that nerve regeneration can be accelerated by increasing G3BP1 acetylation in PNS axons. G3BP1 granules are known to sequester regeneration-associated axonal mRNAs, like *Nrn1* and *Kpnb1*, acting as a barrier to their translation and impeding axon regeneration^4,5^. Rodent PNS nerve injury rapidly stimulates acetylation of axonal G3BP1 on K374 (corresponding to hG3BP1 K376), with the post-translational modification showing peak levels at 3 h post-axotomy. This parallels an increase in axonal G3BP1 S149 phosphorylation following injury^8^. Previous work identified injury-induced CK2α-dependent G3BP1 phosphorylation as a driver for axonal G3BP1 granule disassembly, thereby promoting nerve regeneration^8^. The acetylation site in G3BP1 lies within its RRM domain, and acetylation of hG3BP1^K376^ was shown to release bound mRNAs and cause SG disassembly in U2OS cells^10^. Consistent with this, we show that expressing acetylmimetic G3BP1 selectively increases mRNA translation in axons but not in the soma and accelerates PNS nerve regeneration. This raises the possibility that pathways regulating the acetylation status of G3BP1 constitute previously unrecognized neural repair mechanisms.

Both acetylation and phosphorylation of axonal G3BP1 lead to SG disassembly and both PTMs are rapidly upregulated following nerve injury. It is not clear whether a single G3BP1 molecule can be simultaneously phosphorylated and acetylated or if there is any cross-talk between these post-translational modifications for G3BP1. Cross-talk between lysine acetylation and phosphorylation is known to occur for several different proteins^32^. Phosphorylation can determine subsequent acetylation and vice versa. Further, protein activity can be modulated by dual acetylation and phosphorylation. For example, dual acetylation and phosphorylation of the tumor suppressor p53 after UV exposure triggers cell cycle arrest and/or apoptosis^33^. Assuming the acetylmimetic and phosphomimetic mutants faithfully mimic G3BP1^AcK374^ and G3BP1^PS149^, the lack of increased growth promotion by the hG3BP1^S149E/K376Q^ dual-mimetic mutant compared to the single hG3BP1^K376Q^ mutation argues against cross-talk for simultaneous acetylation and phosphorylation in G3BP1. It is appealing to speculate that different G3BP1 populations are subjected to acetylation vs. phosphorylation. However, our data do not exclude sequential impacts of phosphorylation and acetylation on G3BP1 functions, since the growth promoting effects of the acetylmimetic G3BP1 mutation were lost when an S149A mutation was introduced into the protein.

Previous work has shown that disassembly of axonal G3BP1 granules using the 190-208 G3BP1 CPP or expressing amino acids 141-220 of G3BP1’s IDR1 selectively increases intra-axonal and not soma protein synthesis^3,5^. We see a similar effect with G3BP1 acetylmimetic expression. Further, a non-acetylatable G3BP1 decreased axon growth when endogenous G3BP1 was depleted, a manipulation that was previously shown to increase axon growth^19^. These observations are consistent with non-acetylated G3BP1 sequestering axonal mRNAs away from translation and acetylation of G3BP1 releasing those mRNAs for access to the translational machinery. G3BP1 is a core SG protein and though SG formation has classically represented a response to cellular stress^34^, uninjured PNS axons contain SG-like G3BP1 granules^4,5^. Assuming those axons are not stressed before injury, this indicates that neurons utilize SG-like G3BP1 granules differently than other cell types. However, PNS axons do initiate an intrinsic stress response, with localized release of Ca^2+^ from the endoplasmic reticulum (ER), eIF2α phosphorylation, and translation of stress response mRNAs after axotomy^4,21,35-37^. Beyond neurons, these mechanisms may represent a general strategy through which post-translational modifications of RNA granule proteins fine-tune RNA–protein assemblies, thereby coordinating their transition from translationally silent condensates to active sites of protein synthesis.

Pathological aggregation of G3BP1 with the Tar-DNA binding protein 43 (TDP43) has been demonstrated in PNS motor axons in amyotrophic lateral sclerosis (ALS)^38^. Furthermore, neuronal SG aggregation has been demonstrated in Alzheimer’s disease (AD) brain^39^. Amyloid beta peptide that accumulates in amyloid plaques in AD was shown to stimulate a local stress response that activates translation of *Atf4* mRNA in axons^40^, and dendrites in close proximity to amyloid plaques show elevated Ca^2+^ and alterations in *Camk2α* mRNA translation^41^. These observations linking SGs to neurodegeneration are intriguing given the rapid proximal axon degeneration seen after axotomy in ELP3-depleted PNS neurons that is prevented by acetylmimetic G3BP1. This suggests that stress granules are involved not only in normal axon physiology or stress but also in vulnerability to neurodegeneration.

The enzymes mediating acetylation and deacetylation of G3BP1 were identified in U2OS cells as CREBBP and HDAC6, respectively^10^. Axonal functions for HDAC6 have been established, where inhibition of HDAC6 promotes axon growth on non-permissive substrates and sustains axonal mitochondrial motility^23,24^. Our observations confirm that HDAC6 deacetylates G3BP1 and raise the possibility that axonal G3BP1 granules and sequestered mRNAs are downstream contributors to other functions of axonal HDAC6. Consistent with this, we see that HDAC6 inhibitors also increase axon growth on permissive substrates while decreasing axonal G3BP1 granules and increasing acetylated G3BP1.

Of the ~20 known cytoplasmic KATs in human^42^, we only found ATAT1, CREBBP, and ELP3 in rodent sciatic nerve axoplasm by MS. Despite a report that CREBBP acetylates G3BP1 in U2OS cells^10^, our data indicate that CREBBP depletion does not affect axonal G3BP1 acetylation status. Indeed, axonal G3BP1 granules were actually decreased with CREBBP and ATAT1 depletions. In contrast, ELP3-depletion increased axonal G3BP1 granules, decreased acetylated G3BP1, and decreased axon growth arguing that ELP3 is the axonal acetyltransferase for G3BP1. ELP3 is the catalytic subunit of the Elongator complex, a six-subunit complex that contributes to transcriptional elongation in the nucleus and functions as a KAT in the cytoplasm^43^. ELP3 has been previously shown to acetylate α-Tubulin and histones as well as to post-transcriptionally modify tRNA uridines^27,44–46^. Mutations in ELP3 are causally linked to ALS^18^, and Bento-Abreu *et al*. (2018) more recently implicated loss of ELP3-dependent tRNA modification in motor neuron degeneration seen with ALS^45^. Conditional deletion of murine ELP3 using a ChAT-Cre driver line was very recently shown to result in motor neuron degeneration over 4-14 weeks of life^12^. Decreased expression of *Elp3* mRNA was reported in transcriptome data from human Alzheimer disease (AD) brains and mouse AD models^17^. ELP3 catalyzes 5-methoxycarbonylmethyl (mcm^5^) and 5-carbamoylmethyl (ncm^5^) modifications of the tRNA ‘wobble’ uridine that increases translation fidelity and efficiency^47^. Consequently, decreased ELP3 activity would decrease mRNA translation. Consistent with this, decrease in ELP3 expression in SH-5Y cells expressing amyloid precursor protein with a familial AD mutation show an overall decrease in protein synthesis^17^. The increase in G3BP1 granules that we see with ELP3 depletion would similarly decrease protein synthesis. With the colocalization of G3BP1 with TDP43 aggregates in ALS motor axons^38^, it is intriguing to speculate that increased G3BP1 LLPS occurs with altered ELP3 activity in ALS.

Work in other organisms beyond the mouse neurons studied here indicate that ELP3 plays a role in axon growth. An ethyl methane sulphonate (EMS)-based mutagenesis screen in *Drosophila* identified ELP3 as a critical regulator of axon targeting, and morpholino-based knockdown of *Zebrafish* ELP3 disrupted motor axon growth^18^. Neither these nor other studies addressed what happens to ELP3-depleted axons after injury. Our data surprisingly show that axons proximal to an injury site rapidly degenerate while still connected to the soma. This degeneration is morphologically reminiscent of Wallerian degeneration that occurs in axons separated from the soma^48^. Though ELP3 likely has many substrates in the axons, expression of acetylmimetic hG3BP1 prevented degeneration of the proximal axon segments after injury and fully rescued the axon regeneration phenotype after crush injury in ELP3-depleted neurons. The degeneration seen in proximal ELP3-depleted axons suggests that decreases in axonal ELP3 activity may place neurons at a more vulnerable position in terms of responding to stress.

Together, our findings identify G3BP1 acetylation as a key regulatory mechanism that couples stress granule disassembly to axon regeneration and integrity. We reveal a mechanistic link between axon regeneration and neurodegeneration through the control of stress granule dynamics by post-translationally modified G3BP1. By defining G3BP1 acetylation as both a driver of axonal growth and a safeguard against axotomy-induced degeneration of proximal axons, this work bridges regenerative and degenerative pathways. Targeting the enzymes that regulate G3BP1 acetylation, such as ELP3 and HDAC6, may therefore offer new therapeutic opportunities to fine-tune stress granule dynamics, promote regeneration, and prevent degeneration under chronic stress conditions, including neurodegenerative diseases.

## METHODS

### Animal use

Institutional Animal Care and Use Committees of University of South Carolina and Weizmann Institute of Science approved all animal procedures. Male and female WT (C57BL/6) and G3BP1^FLAG/FLAG^ (both C57BL/6 strain) mice were used for all sciatic nerve injury and DRG cultures. Male Sprague Dawley rats (175–250⍰g) were used for DRG cultures and sciatic nerve axoplasm isolation. Isoflurane was used for anesthesia for AAV transduction and peripheral nerve injuries.

For peripheral nerve injury, anesthetized rats or mice were subjected to a sciatic nerve crush at mid-thigh as described^49,50^. In cases where animals were transduced with AAV prior to injury, viral preparations were injected into the proximal sciatic nerve 14 d prior to crush injury (at sciatic notch level; 1-2⍰×⍰10^12^ or 4-5-2⍰×⍰10^11^ viral particles in 0.6⍰M NaCl).

To isolate axoplasm, 1-1.5 cm of rat or mouse sciatic nerves were collected on ice from niave and 7 day post nerve crush animals. Nerve segments were quickly cut into 2-3 pieces, then placed in nuclear transport buffer without protease inhibitor (20 mM HEPES, 110 mM potassium acetate, and 5 mM magnesium acetate in DEPC-treated water). The axoplasm was isolated by gently squeezing the nerve pieces manually using a pellet pestle in a 1.5 ml microcentrifuge tube. After the axoplasm was manually squeezed out, the samples were centrifuged at 4°C for 10 min at 21,130 xg. The supernatants were then processed for immunoblotting, immunoprecipitation or mass spectrometry (see below).

### Behavioral analyses of nerve regeneration

Behavioral analyses were done in the *Blackbox One* system that uses near-infrared (NIR) light to capture full-body pose, postures, paw placement, gait, rearing, and grooming for mice in a dark enclosed arena. Paw contact and pressure is captured using a proprietary Optitouch sensor, with ‘Frustrated Total Internal Reflection’ (FTIR) illumination generating a ‘paw luminance’ value as a proxy for paw usage. Surface contact by a paw disrupts the internally reflected NIR, creating bright spots quantified as luminance values (0–255) where higher values indicate greater pressure and lower values indicate lighter contact^22^. Thus, denervation of hind limb after sciatic nerve crush results in lighter pressure (i.e., decreased luminance) for the hind paw on the injured side.

Equal numbers of adult male and female C57BL/6 mice were used. All mice were individually housed in standard clear plastic cages under controlled conditions (temperature 21-24°C, humidity 60%, light cycle 12-12) throughout the duration of experiments. For animals subjected to sciatic nerve injections (d 0) + sciatic nerve crush (d 14 post-injection), recordings were done on d 0 prior to injections and then every 3-4 d out to 28 d after the nerve crush (d 42 post-injection). All recordings were done between 10:00 and 17:00 in the same room where the animals were housed using *BlackBox One* instrument (BlackBox Bio, Cambridge, MA US). Animals were placed into individual chambers of the instrument and allowed 10 min of habituation prior to each recording. After 10 min habituation animals were briefly removed to clean the glass surface of urine, feces, and any debris and then returned to individual chambers for 20 min continuous recording.

### Generation of G3BP1^FLAG^ mouse line

For generating the G3BP1^FLAG^ mice, a guide RNA (gRNA) targeting the N-terminus of the G3BP1 gene (Guide: GGTTGAACCCACCAAAGCGA) was designed using a combination of the following tools: the MIT CRISPR design tool^51^ and sgRNA Designer, Rule set 2^52^, in the Benchling implementations (www.benchling.com), SSC^53^ and CRISPOR^54^. A single strand oligonucleotide (ssODN) was used to introduce a 3xFLAG tag (DYKDHDGDYKDHDIDYKDDDDK) and GSG linker. A single proline codon was inserted immediately downstream of the start codon, before the flag tag, in order to disrupt the PAM site, which otherwise would have been recreated by the first codon of the flag tag (ssODN – agtgtagtactgtcgcacaaactcccgcccgaccagcaggggactaggcttctccataactccagatcccttgtcatcgtcatccttgtaatcgatgtcatgatctttataatcaccgtcatggtctttgtagtctggcatcgctttggtgggttcaacCTGTAAAACAATACAAGTCAGTTAACCAATCACAAGAGC).

Cas9 nuclease, crRNA, tracrRNAs and ssODN were purchased from Integrated DNA Technologies (IDT). G3BP1^FLAG^ mice were generated at the Transgenic Facility at the Weizmann Institute of Science using CRISPR/Cas9 genome editing in isolated one-cell mouse embryos as described^55^. C57Bl/6JOlaHsd mice were (Envigo, Israel) maintained in specific pathogen-free condition on a 12 hr light/dark cycle with food and water provided *ad libitum*. Cas9-gRNA ribonucleoprotein complexes together with donor repair template were delivered to one-cell embryos via electroporation, using the Biorad Genepulser. Electroporated embryos were transferred into the oviducts of pseudopregnant ICR females (Envigo, Israel). Genomic DNA from F0 pups was analyzed at weaning for knockin confirmation by PCR and Sanger sequencing with the following primers: Primer1 – TGTTGAGTTGGCTTAGCACAG and Primer 2 - CCTCCTGGGCACTTGAGAG. Mice with verified FLAG-tag were backcrossed at least once to wild-type mice and then bred to homozygosity.

### Cell Culture

For primary neuronal culture, DRGs harvested from adult mice or rats were placed in Hibernate-A medium (BrainBits, HA500), washed 6 times with DMEM/F12 (Corning, 45000-316) and 100 U/ml of Penicillin-Streptomycin (Gibco, 15-140-122) and treated with 0.1 U/µL collagenase type II (Gibco, 17-101-015) 30 min at 37°C, 5% CO2. Ganglia were dissociated into single cell suspensions using a fire-polished Pasteur pipette followed by centrifugation for 10 min at 700 x g. DRGs were washed twice by resuspension with DMEM/F12 (Corning, 45000-316), 10% fetal bovine serum (Gibco, 10437028), and 100 U/ml of Penicillin-Streptomycin (Gibco, 15-140-122) and centrifugation for 10 min at 700 x g. Following washes in fresh medium, cell pellets were resuspended in DMEM/F12 (Corning, 45000-316), 1 x N1 supplement (Sigma, N6530-5ML), 10% fetal bovine serum (Gibco, 10437028), 10⍰μM cytosine arabinoside (Sigma, C1768-500MG), 2 mM L-glutamine (Gibco, A2916801), and 100 U/mL of Penicillin-Streptomycin (Gibco, 15-140-122). Resuspended DRGs were transfected or plated immediately on laminin/poly-L-lysine-coated substrates. For coating substrates, coverslips or dishes were incubated in 50 µg/ml poly-L-lysine (Fisher, A005C) overnight at 37°C and 5 µg/ml laminin (Millipore, CC095-5MG) for 4 h at 37°C. #1.5 glass coverslips 12 mm round (Warner Instruments, 64-0712), 35 mm black wall glass bottom dishes (WillCo Wells from Ted Paella, HBSB-3522), or 6 well plastic dishes were used for culture (Corning, 3516). Culture medium was changed the day after plating. After 60-65 h, cultures were rinsed with 1X PBS and processed for immunoblotting and immunoprecipitation or fixed with 4 % paraformaldehyde and processed for immunofluorescence or PLA (see below).

For spot cultures, mouse DRGs were collected and placed in Hibernate-A medium (BrainBits, HA500) on ice. DRGs were washed six times with DMEM/F12 (Corning, 16-405-CV), 10% fetal bovine serum (Gibco, 10437028) and 100 U/ml of Penicillin-Streptomycin (Gibco, 15-140-122) and treated with 0.1 U/µl collagenase type II (Gibco, 17-101-015) for 15 min at 37°C, 5% CO2. Following incubation, the cells were gently triturated 10 times using a 1⍰ml pipette, then returned to the incubator for an additional 10⍰mins at 37°C, 5 %CO_2_. The cells were triturated again 10 times with a 1 ml pipette, transferred to a 15⍰ml tube, and washed twice with culture medium. In between washes, cells were centrifuged at 700 x g. In parallel, laminin-coated plates were rinsed twice with 1X PBS and allowed to dry completely under the hood without adding any medium. For coating substrates, 4 well chambers were incubated in 1X poly-D-lysine (Fisher, P7280) overnight at 37°C and 5 µg/ml laminin (Millipore, CC095) for 4 h at 37°C. Cells were resuspended in culture medium according to the required plating density; typically, DRGs from one mouse (35–40 DRGs) were resuspended in 56⍰µL of DMEM/F12 (Corning, 45000-316), 1X N21 supplement (R&D Systems, AR008), 10% fetal bovine serum (Gibco, 10437028), 10⍰μM cytosine arabinoside (Sigma, C1768-500MG), 1X GlutaMAX™ Supplement (Gibco, 35050061), and 100 U/ml of Penicillin-Streptomycin (Gibco, 15-140-122) to prepare eight spots (7⍰µl per spot). Using a 20⍰µl pipette, 7⍰µl of the resuspended cells was carefully plated onto each spot on the 4 well chambers (Ibidi, 80297), 2 spots per chamber. The cultures were returned to the incubator without additional medium for 7⍰min. Culture medium (as above) was added carefully to avoid disturbing the spots, and the plates were returned to the incubator. Spot cultures were incubated at 37°C at 37°C, 5% CO_2_ for 7-8 d. For media changes, half volume of the medium in each well was removed and replenished with fresh DRG culture medium at 1 and 5 d. For axotomy, axons were cut on one side of the spot under an inverted microscope using a sterilize 0.6 mm thick x 2.75 mm wide flat carbon steel surgical blade (Fine Science Tools, 10035-10)^29^. Axon regrowth or degeneration from the injury site was evaluated every 10 min for up to 16 h post-axotomy or as indicated.

For viral transduction in spot cultures, dissociated DRGs were resuspended in 52 µl culture medium as above and 4 µl of 4.14 x 10^11^ GC/ml AAV-PHPS-shElp3 or AAV-PHPS-shCntl were added per 8 spots. For the rescue experiment, 2 µl of 4.14 x 10^11^ GC/ml of AAV-PHPS-shElp3 or AAV-PHPS-shCntl and 2 µl of 1.39 x 10^12^ GC/ml AAV9-mCherry-G3BP1^K376Q^ or AAV9-mcherry were added and DRGs were then incubated in ice for 5 min before plating. After putting 7 µl of DRGs mixed with the virus for each spot, plates were put in the incubator for 8 min. Finally 700 µl of culture media were delicately added to not disturb the spots.

For cDNA transfection, dissociated ganglia were pelleted by centrifugation at 700 x g for 5⍰min and resuspended in ‘nucleofector solution’ (Rat Neuron Nucleofector kit; Lonza, VSPI-1003). 5-7⍰μg plasmid was electroporated using an AMAXA Nucleofector apparatus (program G-013). Transfected DRGs were plated as above.

For siRNA transfection, 25⍰nM siRNAs (Horizon) were used with DharmaFECT 3 (Horizon, T-2003-02) reagent and incubated for 72⍰h. Three *G3bp1* 3’UTR targets siRNAs were used to deplete G3BP1 (5’ CCCAUAGGAGCUGGGAAUUU 3’, 5’ GUAGUAAAGUGUAUGGUUAUU 3’ and 5’ GGGAAGAGAUGGUGGGAAGUU 3’). siRNAs for *Elp3, Crebbp1* (CBP/P300) and *Atat1* mRNAs were used to deplete ELP3, CREBBP1 and ATAT1 (Horizon L-045781-01-0005, L-050973-02-0005, and L-041130-00-0005, respectively). Non-targeting siRNAs were used as control in each experiment (Horizon, D-001810-10). RTddPCR were used to test the efficiency of *G3bp1, Elp3, Crebbp1* and *Atat1* depletion (see below).

For Tubastatin A and NN-769 treatment, compounds were diluted in growth media and 10 µM TubA was bath applied to dissociated DRGs and then cells lysed and processed for immunoblotting (see below). 10 µM (TubA) or 10 nM (NN-769) was bath applied to dissociated DRGs for 2h for axonal growth analysis and immunostaining. Equal volume of DMSO was used as vehicle control.

### Plasmid and viral expression constructs

hG3BP1-mCh, hG3BP1^K376Q^-mCh, hG3BP1^K376R^-mCh were previously published^10^. mCh fusions for hG3BP1^S149A^ and hG3BP1^S149E^ were generated by cloning from previously published BFP fusion constructs^8^. hG3BP1^S149A^-mCh, hG3BP1^S149E^-mCh, hG3BP1^K376Q/S149A^-mCh, hG3BP1^K376R/S149A^-mCh, hG3BP1^K376Q/S149E^-mCh, and hG3BP1^K376R/S149E^-mCh were generated by restriction enzyme cloning using BsrG1 and *Kpn1*, with hG3BP1-mCh, hG3BP1^K376Q^-mCh, and hG3BP1^K376R^-mCh as backbones. All plasmid inserts were fully sequenced prior to use.

AAV preparations were designed and generated by *Vectorbuilder*. AAV-PHPS-BFP-shElp3 (pAAV[shRNA]-CAG>EBFP-U6>elp3_shRNA; target region 304-324 nt in NCBI Gene ID # 74195) and AAV-PHPS-GFP-shCntl (pAAV[shRNA]-CAG>EGFP-U6>Scramble_shRNA (cat # AAV9M VB010000-9489hhg).

### RNA isolation and ddPCR analyses

RNA was isolated from dissociated DRG cultures using the RNeasy Microisolation kit (Qiagen, 74104). Fluorimetry with Quant-iT RiboGreen RNA Reagent (ThermoFisher, R11491) was used for RNA quantification. For analyses of total RNA levels, RNA yields were normalized across samples and then reverse transcribed using Sensifast cDNA synthesis kit per manufacturer instructions (Thomas Scientific, C755H66). ddPCR products were detected using QX200 ddPCR EvaGreen Supermix (Biorad, 1864034) and QX200TM droplet reader (Biorad). *Gapdh* and *Hmgb1* mRNAs signals were used for normalizing yields across different isolates as indicated in results. The following primers were used for ddPCR (all from Integrated DNA Technologies): *G3bp1* forward 5’-CCTTTGTCCTTGCTCCTGA-3’ and reverse 5’-CTTCTACTTCTTCCTCGGATTCC-3’; *Elp3* forward 5’-GGAGAGAGACATTGAGCAGTT-3’ and reverse 5’-GCTCATACAGACCAGTTCCAC-3’; *Crebbp1* forward 5’-TCACAATCAACATCTCCTTCCC-3’ and reverse 5’-TGTCGATAGAGTGCTTCTAGAGT-3’; *Atat1* forward 5’-TCCTGAATAAGCACTACAACCTG-3’ and reverse 5’-GAGTGTCGAGTTGCTCTCAG-3’; *Gapdh* forward 5’-AATGGTGAAGGTCGGTGTG-3’, reverse 5’-GTGGAGTCATACTGGAACATGTAG-3’; and, *Hmgb1* forward 5’-TGGCAAAGCAAGGAGTGAT-3’ and reverse 5’-AATGGCGGTTAAAGGAGAGTC-3’.

### Nascent protein synthesis detection

To visualize newly synthesized proteins in cultured neurons, we used the Click-it™ Plus OPP Alexa Fluor™ 488 Protein Synthesis Assay Kit per manufacturer’s instructions (Invitrogen/Life Technologies). 3 d after transfection, rat DRG cultures were incubated with 20 µM O-propargyl-puromycin (OPP) for 30⍰min at 37°C. OPP-labeled proteins were detected by crosslinking with Alexa Fluor-488 picolyl azide molecule. Coverslips were then rinsed in PBS, fixed with buffered 4% PFA and mounted with Prolong Gold Antifade (Invitrogen) and imaged with Leica DMI6000 epifluorescent microscope. FIJI/ImageJ was used to quantify the Puromycinylation signals in the axon shaft (≥⍰200⍰µm from cell body) and cell bodies.

### Immunofluorescence (IF)

All procedures were performed at room temperature unless specified otherwise. Cultured neurons were fixed in buffered 4% PFA for 15 min. Neurons were permeabilized with 0.3% Triton X-100 in PBS for 5 min and incubated for 1 h in blocking buffer (0.1% Triton X-100 or 0.1% Tween + PBS and 10% Normal Donkey Serum (Jackson ImmunoResearch, 017-000-121). Neurons were then incubated with primary antibodies diluted in blocking buffer overnight in a humidified chambers at 4°C.

For IF in tissues, sciatic nerves were fixed overnight in buffered 4% PFA, followed by cryoprotection for 2-3 days in buffered 30 % sucrose, both at 4°C. Nerves were embedded in OCT and processed for cryosectioning at 12⍰µm thickness. Sections were washed in buffered 1x PBS, followed 10 mM glycine 3 times for 10 min each and 0.25 M NaBH_4_ 3 times for 10 min each. Sections were permeabilized for 15 min in 0.3% Triton X-100 in 1X PBS, washed in 1X PBS 3 times, 5 min each, and then blocked for 1 h in 1X PBS containing 10% normal donkey serum (Jackson ImmunoResearch, 017-000-121) + 0.1% Triton-X. After blocking, sections were incubated overnight in primary antibodies diluted in blocking buffer at 4°C in a humidified chamber. Sections were washed three times in 1X PBS and then incubated with secondary antibodies diluted in blocking buffer for 1 h. Samples were washed in 1X PBS three times, briefly rinsed in deionized water and mounted with Prolong Gold Antifade with or without DAPI (Invitrogen, P36931 and P36930).

Primary antibodies used for immunofluorescence staining on tissues consisted of: chicken anti-beta III tubulin (1:100, Millipore, AB9354), chicken NF 200 kDa (1:100, Aves, NFH88697979), chicken NF 160 kDa (1:100, Aves, NFM8887979), chicken NF 68 kDa (1:100, Aves, NFL87817983), rabbit anti-G3BP1 (1:200, Sigma, HPA004052), rabbit anti-G3BP1^AcK374^ (1:100)^10^, rabbit anti-SCG10 (Stathmin-2, 1:100, Novus Biologicals, NBP1-49461), RT97 mouse anti-NF (NF; 1:200, Devel. Studies Hybridoma Bank, AB_528399), SMI312 mouse anti-neurofilament (1:200, BioLegend, 837904), TUJ mouse anti-tubulin beta III (1:200, BioLegend, 801201), chicken anti-GFP (1:200, Abcam, ab13970), mouse Ac-Lys monoclonal antibody (1C6) (1:100 Thermofisher, MA1-2021), Mouse DegenoTag™ (1:1000, EnCor, MCA-1D44), and mouse Anti-RFP mAb (1:200, MBL, M155-3). Secondary antibodies used for immunofluorescence staining on tissues consisted of: DyLight 405-conjugated donkey anti-chicken (703-475-155), FITC-conjugated donkey anti-chicken (703-095-155), FITC-conjugated donkey anti-rabbit (711-095-152), FITC-conjugated donkey anti-mouse (715-095-150), Cy3-conjugated donkey anti-chicken (703-165-155), Cy3-conjugated donkey anti-rabbit (711-165-152), Cy 3-conjugated donkey anti-mouse (715-165-150), Cy5-conjugated donkey anti-chicken (703-175-155), and Cy5-conjugated donkey anti-rabbit (711-175-152; all from Jackson ImmunoResearch, used at 1:200 for tissue section).

Primary antibodies used for immunofluorescence staining on DRGs culture consisted of: chicken anti-beta III tubulin (1:200), chicken NF 200, 160 and 68 kDa (1:200), rabbit anti-G3BP1 (1:200), rabbit anti-G3BP1^AcK374^ (1:300)^10^, RT97 mouse anti-NF (NF; 1:200), SMI312 mouse anti-neurofilament (1:200), purified TUJ mouse anti-tubulin beta III (1:200), mouse anti-FLAG (1:500), 1C6 mouse Ac-Lys monoclonal antibody (1:100) and rabbit Anti-KAT9/Elp3 antibody (1:200, Abcam, ab190907). Secondary antibodies used for immunofluorescence staining on DRGs cultures consisted of: DyLight 405-conjugated donkey anti-chicken, FITC-conjugated donkey anti-chicken, FITC-conjugated donkey anti-rabbit, FITC-conjugated donkey anti-mouse, Cy3-conjugated donkey anti-chicken, Cy3-conjugated donkey anti-rabbit, Cy 3-conjugated donkey anti-mouse, Cy5-conjugated donkey anti-chicken, Cy5-conjugated donkey anti-rabbit, and Cy5-conjugated donkey anti-mouse (all from Jackson ImmunoResearch, used at 1:400).

### Proximity Ligation Assays (PLA)

NaveniFlex Cell Atto647N kit (Navinci, 60017) was used for PLA per the manufacturer’s instructions. Briefly, cells were fixed with buffered 4% PFA for 15 min and treated with 0.3 % Triton for 5 min. DRGs were then blocked in 0.1% Tween in PBS containing 10% normal donkey serum (Gibco, 10437028) for 1 h at 37°C and primary antibodies were applied diluted in the blocking solution for 1 h at 37°C (mouse anti-FLAG (1:500); rabbit Ac-Lys (1:100, Cell Signaling, 9441)). Navenibody M1 and Navenibody M2 were incubated for 1 h at 37°C followed by Naveniflex reaction 1 and reaction 2. Finally, cells were post fixed with buffered 4% PFA for 15 min and immunofluorestaining for NF was performed as above (chicken NF 200, 160 and 68 kDa (1:200 each). Coverslips were mounted on glass slides using Prolong Gold Antifade.

### Immunoblotting

Adult rat DRGs (plated in 6 well plate at 20 DRGs/well) were rinsed with buffered 1X PBS twice, lysed in RIPA buffer (50 mM Tris-HCL, 150 mM NaCl, 1% NP40, 0.1% SDS, and 0.5% sodium-deoxycholate with 1 x protease and phosphatase inhibitors (Thermo Scientific, PIA32965 and PIA32957) plus 10 µM TubA), and then passed through a 30 Ga syringe 6 times. Following centrifugation to clear the lysate at 12000 x g for 20 min at 4°C, protein concentration of the supernatant was determined using BCA assay (Pierce, 23227). Lysates were normalized for protein content, denatured by boiling for 5 min in 1x Laemmli buffer, fractionated by SDS-PAGE, and transferred to PVDF membrane.

Blots were blocked for 1⍰h at room temperature with 3% BSA in Tris-buffered saline with 0.1% Tween 20 (TBS-T). Primary antibodies diluted in blocking buffer were added to the membranes and incubated overnight at 4°C with rocking. Primary antibodies consisted of: rabbit anti-G3BP1 (1:2000, Sigma, HPA004052), rabbit anti-GAPDH (1:2000, Cell Signaling, 5174S), and rabbit anti-G3BP1^AcK374^ (1:500)^10^. After washing in TBS-T, blots were incubated with HRP-conjugated anti-rabbit IgG antibodies (1:5000, Cell Signaling 7074) diluted in blocking buffer for 1⍰h at room temperature. After washing in TBS-T, signals were detected by chemiluminescence using ECL Prime^TM^ (GE Healthcare, GERPN2236) or Clarity Max Western ECL Substrate (Biorad, AM9759). Mouse Monoclonal ANTI-FLAG® M2-Peroxidase (HRP) antibody (1:1000, Sigma, A8592) was used for detection of G3BP1^FLAG^, so those blots were processed for chemiluminescence as above immediately following primary antibody incubation and wash.

### Immunoprecipitations (IP)

Immediately after dissection, six mouse sciatic nerve (2-3 cm segments) were placed into two separate 1.5 ml tubes containing 200 µl nuclear transport buffer (20⍰mM HEPES [pH 7.3], 110⍰mM potassium acetate, and 5⍰mM magnesium acetate with 1 x protease and phosphatase inhibitors). Nerves were then cut into 1-2 mm pieces and the axoplasm was gently extruded using a pestle. Nerves were centrifuged at 12,000 x g for 20 min at 4°C. Equal volume of 2 x RIPA was added to the supernatant and an aliquot was used for quantification with Pierce™ BCA Protein Assay Kit (Pierce, 23227). After normalizing the yields, 10% of axoplasm was saved as input. The remainder was precleared with 30 µl of RIPA buffer equilibrated Dynabeads™ Protein A for Immunoprecipitation (Invitrogen, 10002D) rotating for 30 min at 4°C. Axoplasm was then incubated with 40 µl of pre-cleared in RIPA buffer Anti-FLAG® M2 Magnetic Beads overnight at 4°C with rotation. After 6 washes in RIPA Buffer, FLAG-G3BP1 was eluted using 1x SDS Laemmli buffer and processed for immunoblotting as above.

### Live cell imaging

Spot culture neurons that had been transduced were imaged after 7 DIV. Axons were transected on the inverted microscope platform of a Leica *Stellaris* confocal microscope. Images were then acquired by confocal microscope with environmental chamber maintained at 37°C, 5% CO_2_ using a 63⍰× ⍰/1.4 NA oil immersion objective. Signals were imaged as single optical planes or Z-stacks every 10 min for 16 h (AAV-PHPS-BFP-shElp3: 405⍰nm excitation and 8.78 % white light laser power; 430-568 nm emission; AAV-PHPS-GFP-shCntl: 485⍰nm excitation and 21.08 % white light laser power; 490–750⍰nm emission).

### Stimulated emission depletion microscopy (STED)

Leica Stellaris confocal microscope with HyD detectors was used for imaging. Signals were imaged as single optical planes (for G3BP1, 554 nm excitation and 2 % white light laser power, 559-631 nm emission, and 660 nm depletion laser; for Ac-Lys 490 nm excitation and 2 % white light laser power, 495-553 nm emission, 594 depletion laser). Primary antibodies were: rabbit anti-G3BP1 (1:100), 1C6 mouse Ac-Lys antibody (1:100) and a cocktail of chicken NF 200, 160 kDa and 68 kDa (at 1:200 each). Secondary antibodies were: donkey anti-mouse AF488, Cy3-conjugated donkey anti-rabbit, and Cy5-conjugated donkey anti-chicken (all at 1:200).

### Mass spectrometry

Naïve and regenerating sciatic nerve axoplasm from 6 rats that had undergone unilateral nerve crush 7 days before harvest was isolated as above. 900 µl of Trizol was added to the extruded axoplasm and then the samples were kept frozen until processing. Samples were thawed and supplemented with 180 µl of chloroform, shaken vigorously for 30 sec and incubated at RT for 2-3 min. The resulting mixture was centrifuged at 12,000 xg for 15 min at 4 °C. The clear aqueous phase overlaying the interphase was carefully removed and discarded. The remaining fractions (interphase and organic phase), were supplemented with 270 µl ethanol, mixed by inversion. After standing for 2-3 min at room temperature, samples were centrifuged at 2,000 xg for 5 min at 2-8°C. Proteins were precipitated from the supernatant by mixing with 1.35 ml 2-propanol, incubated 10 min at room temperature, followed by centrifugation at 12,000 xg for 10 min at 2–8 °C. The resulting protein pellet was washed by adding 1 ml of 0.3 M guanidine hydrochloride in 95% ethanol, incubating for 20 min at room temperature, and then removing supernatant. Protein pellet was washed 2 more times with 1 ml 0.3 M guanidine hydrochloride in 95% ethanol. After the final wash, the supernatant was removed and 1 mL 100% EtOH was added to each pellet, vortexed and incubated 20 min at RT. After centrifuging (12,000 xg for 10 min) the supernatant was removed and the protein pellets dried with Spin-Vac for 5-10 min. Pellets were resuspended in 50 µl 8M guanidine hydrochloridel and sonicated in a bath sonicator for 15 min. Protein concentration was measured using the Pierce™ BCA Protein Assay Kit (Thermo Scientific) in triplicates, taking 1 µl aliquots of this solution. Aliquots of samples containing 300 µg of protein were added to nine fold the sample volume of ethanol and allowed to precipitate at −80°C overnight. Pellets were centrifuged at 15000 xg 10 min, supernatant removed, and the samples were allowed to dry. The pellets were resuspended in 12.5 µl 6.25 M guanidine hydrochloride containing 6.25 % (V/V) Sigma-Aldrich Phosphatase inhibitors cocktails 2 and 3, plus 78 mM triethylammonium bicarbonate (TEAB) pH 8.0, and 7.8 mM Tris(2-carboxyethyl)phosphine hydrochloride. Samples were sonicated in a bath sonicator for 5 min, then incubated at 56°C for 15 min, followed by a 30 min incubation at room temperature in the dark with 20 mM iodoacetamide. The samples were added 62.5 µl TEAB containing 2 µg Lysyl endopeptidase (LysC) (FUJIFILM-Wako), and incubated at 37°C overnight. After that, the samples were added 5% (W/W) modified trypsin (TPCK-Trypsin, Thermo Scientific), and incubated overnight at 37°C.

Digested Samples were labeled according to *TMTPro-18* plex kit instructions (ThermoFisher), with some modifications. Briefly, TMT reagents were dissolved in acetonitrile at 25 µg/µl, and 20 µl of these stocks added to the samples (500 µg reagent, 1.7 fold the peptide mass amount). After incubation for 1 h at room temperature samples were quenched with 2 µl 5% hydroxylamine, then all samples were combined adding them over 20 ml 0.1% formic acid, and desalted using a C18 SepPak cartridge (Waters). The Sep Pak eluate was dried in preparation for high pH reverse phase chromatographic fractionation.

The sample was fractionated on an AKTA purifier system utilizing a Phenomenex Gemini 5 u C18 110A 150 x 4.60 mm column, operating at a flow rate of 0.550 ml/min. Buffer A consisted of 20 mM ammonium formate (pH 10), and buffer B consisted of 20 mM ammonium formate in 90% acetonitrile (pH 10). Gradient details were as follows: 1 % to 30% B in 49 min, 30% B to 70% B in 4 min, 70% B down to 1% B in 4 min. Peptide-containing fractions were collected, evaporated and resuspended in 0.1% formic for LC-MS/MS analysis.

For peptide and protein identification and TMT quantitation, aliquots (containing around 3 µg of digested material) coming from the above reverse phase fractionation were run onto a 2 μm, 75 μm ID x 50 cm PepMap RSLC C18 EasySpray column (Thermo Scientific). 3 h MeCN gradients (2-25% in 0.1% formic acid) were used to separate peptides, at a flow rate of 200 nl/min, for analysis in a Orbitrap Exploris 480 (Thermo Scientific) in positive ion mode with the following settings. MS spectra were acquired between 375 and 1,500 m/z with a resolution of 12,0000. For each MS spectrum, multiply charged ions over the selected threshold (2E4) were selected for MS/MS in cycles of 3 s with an isolation window of 0.7 m/z. Precursor ions were fragmented by HCD using stepped relative collision energies of 30, 35 and 45 in order to ensure efficient generation of sequence ions as well as TMT reporter ions. MS/MS spectra were acquired in centroid mode with resolution 60,000 from m/z=120. A dynamic exclusion window was applied which prevented the same m/z from being selected for 30s after its acquisition. Peak lists were generated using PAVA in-house software^56^. All generated peak lists were searched against the *Rattus norvegicus* subset of the UniProtKB database (UniProtKB.2017.11.01), using Protein Prospector^57^ with the following parameters: Enzyme specificity was set as Trypsin, and up to 2 missed cleavages per peptide were allowed. Carbamidomethylation of cysteine residues, and TMTPro16plex labeling of lysine residues and N-termini of the peptides were allowed as fixed modifications. N-acetylation of the N-terminus of the protein, loss of protein N-terminal methionine, pyroglutamate formation from peptide N-terminal glutamines, and oxidation of methionine were allowed as variable modifications Mass tolerance was 5 ppm in MS and 30 ppm in MS/MS. The false discovery rate was estimated by searching the data using a concatenated database which contains the original UniProKB database, as well as a version of each original entry where the sequence has been randomized. A 1% FDR was permitted at the protein and peptide level. For quantitation only unique peptides were considered; peptides common to several proteins were not used for quantitative analysis. Relative quantization of peptide abundance was performed via calculation of the intensity of reporter ions corresponding to the different TMT labels, present in MS/MS spectra. Intensities were determined by Protein Prospector. Summed intensity per sample on each TMT channel for all identified spectra were used to normalize individual intensity values. Relative abundances were calculated as ratios vs the average intensity levels in the channels corresponding to the relevant control samples. Spectra representing replicate measurements of the same peptide were kept. Individual spectral ratios were aggregated to the peptide level using median values of the log_2_ ratios. For total protein relative levels, peptide ratios were aggregated to the protein levels using median values of the log_2_ ratios. Statistical significance was calculated at the protein level by comparing the values for the 3 biological replicates of the relevant groups in the TMT experiment with a 2-tailed t-test.

### Image analyses

For quantitation between samples, imaging parameters were matched for exposure, gain, offset and post-processing.

For analyses of protein levels in tissues, z planes of the xyz tile scans from 3-5 locations along each nerve section were analyzed using FIJI/ImageJ. Colocalization plug-in was used to extract protein signals that overlap with axonal marker (NF) in each plane, with the extracted ‘axon-only’ signal projected as a separate channel.

For automated quantification of axon growth morphologies in cultured neurons, images from 60-65⍰h DRG cultures were taken using the high content image screening ImageXpress micro (Molecular Devices) and analyzed for neurite outgrowth using *WIS-NeuroMath* software tool^58^.

For OPP overall protein synthesis quantification, mCherry-G3BP1, mCherry-G3BP1^K376Q^, and mCherry-G3BP1^K376R^ transfected axons axon shaft (≥⍰200⍰µm from cell body) and cell bodies were considered. Absolute signal intensity was quantified for axonal or cell body only signal using FIJI/ImageJ.

To assess regeneration *in vivo*, tile scans of SCG10-stained nerve sections were post-processed by Straighten plug-in for FIJI/ImageJ. SCG10 positive axon profiles were then counted in 100⍰µm bins at 0.4 or 0.5⍰mm intervals distal from crush site. Fluorescent intensity of axon profiles present in the proximal crush site was treated as the baseline, and values from the distal bins were normalized to this to calculate the percentage of regenerating axons. For regeneration analysis of transduced axons, SCG10 and virus signal were colocalized using ImageJ colocalization plugin. The colocalize axon profiles were then analyzed.

For analysis of Proximity Ligation Assay (PLA) images or G3BP1 granules analysis, 100 µm of axons axon shaft (⍰≥⍰200⍰µm from cell body) was considered. Thresholding was applied to acquired image sequences using FIJI/ImageJ to generate binary masks and FIJI/ImageJ particle analyzer was used to obtain data on number and size of G3BP1 granules.

For analysis of protein levels in DRGs cultures, xy image were capture of 100⍰µm segments of the axon shaft (separated from the cell body and growth cone by⍰≥ ⍰200⍰µm) and analyze for fluorescent intensity using FIJI/ImageJ. Protein signal intensities were then normalized to NF immunoreactivity area.

For degeneration index calculation, 140-150 µm of axon shaft proximal to the transection site were analyzed. Briefly, time-lapse videos were uploaded on ImageJ and thresholded using the “Triangle” option (equivalent parameters were used for thresholding between conditions). For each time point the area occupied by the axons was calculated using ImageJ. Particle analyzer was then used to obtain the area occupied by the fragmented axons (1-3 µm^2^). The degeneration index was then calculated by dividing the area of the fragmented axons over the total axon area. The same analysis was used to calculate the degeneration index for the axons co-transduced with both AAV-PHPS-shElp3 or AAV-PHPS-shCntl plus AAV9-mCherry-G3BP1^K376Q^ or AAV9-mcherry. Co-transduced axons were segmented using FIJI/ImageJ colocalization plugin prior to fragment analyses.

For degeneration quantification of DegenoTag signals in sciatic nerve section, DegenoTag positive axon profiles were counted in 100⍰µm bins at 0.2 mm proximal to the crush site (i.e., towards the DRGs and spinal cord) and 1 and 2 mm distal to the crush site.

### Statistical analyses

*Prism version 7* (GraphPad) and *Excel* (Microsoft) software packages were used for statistical analyses. Two-way or one-way ANOVA was used to compare means of independent groups with indicated post-hoc test and either Student’s t-test or Mann-Whitney was used based on normalization test to compare smaller sample sizes of the in vitro analyses. p values of⍰≤ ⍰0.05 were considered as statistically significant.

## AUTHOR CONTRIBUTIONS

Conceptualization – IDC, JYL, NS, SBD, EP, PKS, and JLT

Data curation – IDC, EM, TS, and JOP

Formal analysis – IDC, EM, SS, AT, JOP, and MM

Funding acquisition – MF, HZ, ALB, EP, and JLT

Investigation – IDC, EM, SS, AT, CNB, JYL, JOP, CM, ET, NS, SBD, RHK, MM, and TS

Methodology – IDC, JYL, SBD, RHK, PKS, JOP, and TS

Resources – CM, NS, SBD, RHK, NN, PM, PTG, and MF

Project administration – ET, HZ, MF, ALB, and JLT

Supervision – JOP, MF, ALB, HZ, and JLT

Validation – IDC, EM, SS, CBN, and JYL

Visualization – IDC, EM, and JLT

Writing, original draft – IDC and JLT

Writing, review & editing – all

All authors approved the submitted version of the paper and have agreed to be personally accountable for their own contributions.

## ACKNOWELDGEMENTS

JL Twiss is the incipient SC SmartState Chair in Childhood Neurotherapeutics at the University of South Carolina.

## FUNDING

Grants from the following sources funded this work: National Institutes of Health (R01-NS117821 and R01-NS089633 to JLT; R01-NS115507 and R21NS-122169 to HZ), Department of Veteran Affairs (01BX002149 and IK6BX006316 to HZ), Dr. Miriam and Sheldon G. Adelson Medical Research Foundation (to ALB, MF and JLT), Merkin Peripheral Neuropathy and Nerve Regeneration Center (to PKS), Canadian Institutes of Health Research and Infrastructure grants from the Canadian Foundation for Innovation and Ontario Research Fund (PTG) and Mitacs through e-Accelerate Program (PM and NN).

## DECLARATION OF INTERESTS

N.N., P.M., and P.T.G. Are cofounders of HDAX Therapeutics.

## DECLARATION OF AI AND AI-ASSISTIVE TECHNOLOGIES

Not applicable.

## SUPPLEMENTAL MATERIALS

### SUPPLEMENTARY FIGURE LEGENDS

**Supplementary Figure 1:**
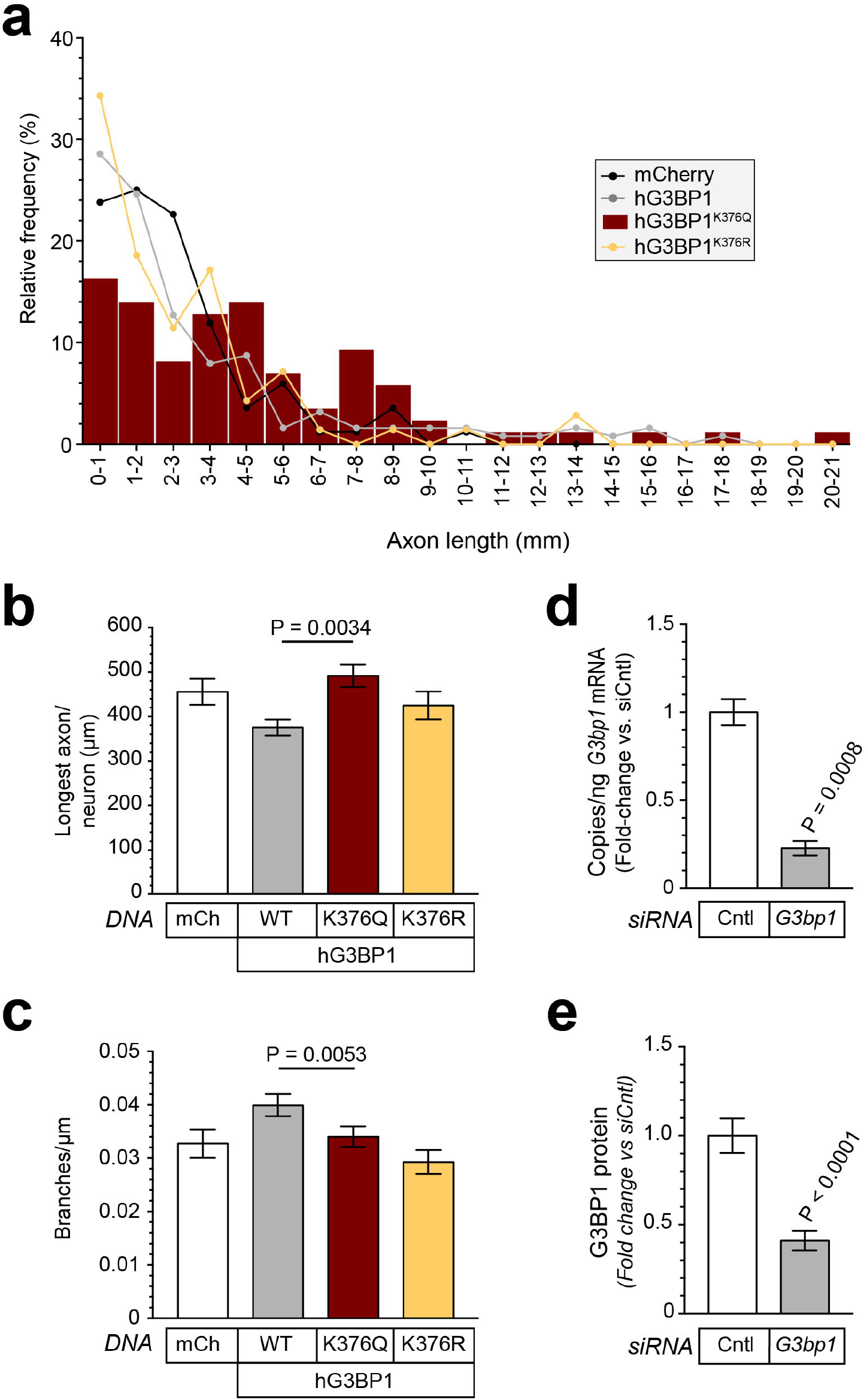
Acetylmimetic hG3BP1 affects axonal growth. **a)**, Histogram comparing axon length distributions in DRG neurons transfected with mCh, WT hG3BP1, and hG3BP1 mutants from Figure 2b-c (1000 µm bins). **b-c)**, Longest axon per neuron (**b**) and axon branching (**c**) for DRGs transfected with mCh, WT hG3BP1, and hG3BP1 mutants from Figure 2c. **d-e)**, *G3bp1* mRNA (**d**) and G3BP1 protein (**e**) levels in DRGs transfected with siCntl vs. siG3BP1 (N ≥ 100 axons per condition for a-c and 115 for e across 3 replicate cultures; 3 replicate cultures for RTddPCR in d; P values by one-way ANOVA with Tukey’s post hoc in b-c and students T-test in d-e).

**Supplementary Figure 2:**
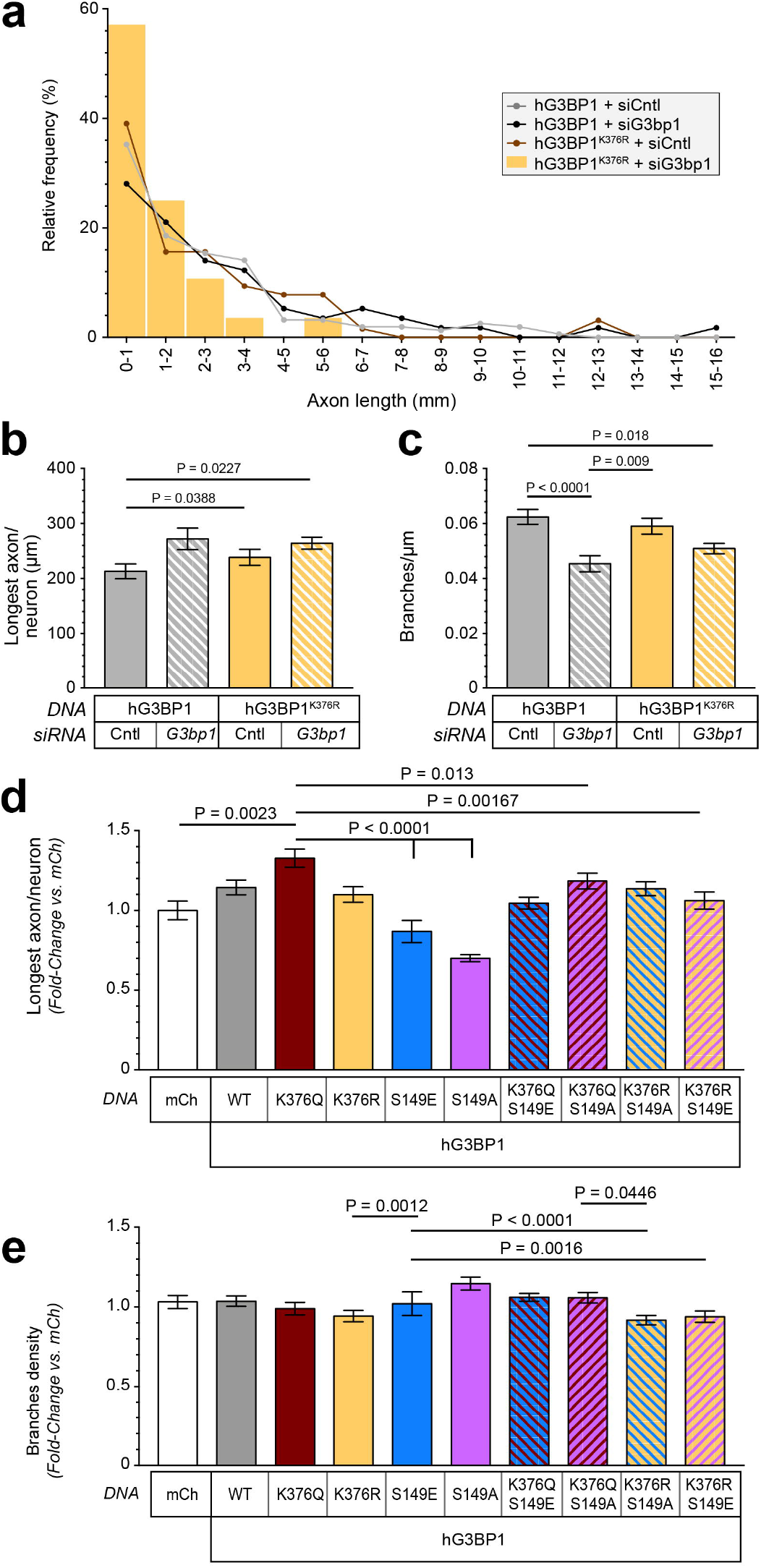
Acetylmimetic and phosphomimetic G3BP1 double mutants effects on axon growth. **a)**, Histogram comparing axon length distributions in DRG neurons cotransfected with WT hG3BP1 or hG3BP1^K376Q^ and siCntl or siG3bp1 from Figure 2e-f (1000 µm bins). **b-c)**, Longest axon per neuron (**b**) and axon branching (**c**) for DRGs transfected with mCh and indicated hG3BP1 constructs plus siCntl or siG3bp1 from Figure 2e-f. **d-e)**, Longest axon per neuron (**d**) and axon branching (**e**) for DRGs transfected with mCh, hG3BP1 WT, and indicated single and double hG3BP1 mutants from Figure 2g (N ≥ 52 neurons per condition across 3 replicated cultures in a-c and ≥ 102 in d-e; P values by one-way ANOVA with Tukey’s post-hoc in b-e).

**Supplementary Figure 3:**
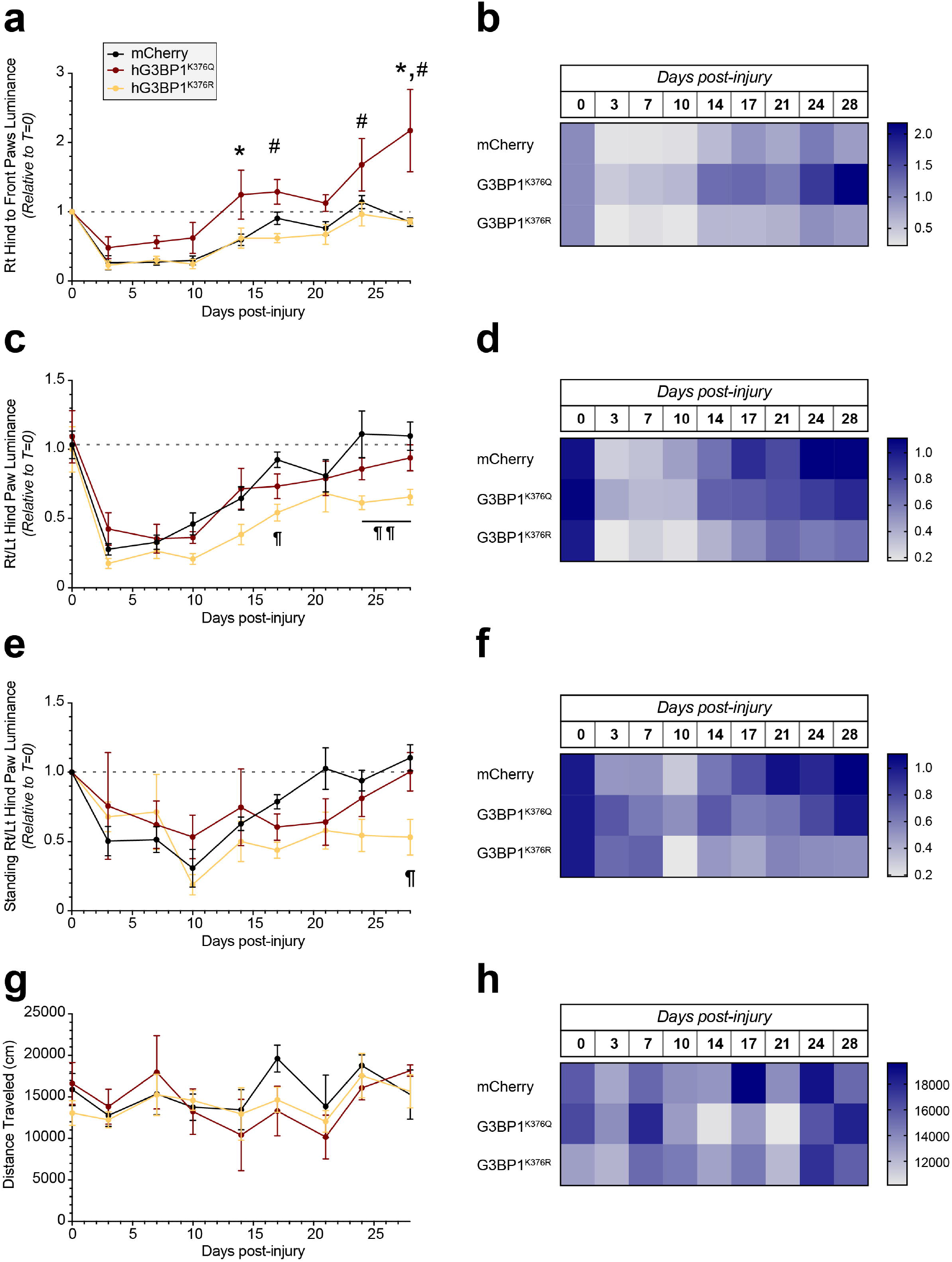
hG3BP1^K376Q^ accelerates nerve regeneration. Functional parameters assess by BlackBox One instrument for AAV-mCh, -hG3BP1^K376Q^ and -hG3BP1^K376R^ transduced mice are shown for indicated times after right (Rt)-sided sciatic nerve crush injury (bilateral AAV transduction at 14 d prior to nerve crush injury). The left side (Lt) underwent sham procedure where nerve was exposed but not further manipulated. Mean ± SEM (a, c, e, and g) and heat maps (b, d, f, and h) are shown for Rt hind paw to average front paws luminance (a,b), Rt to Lt hind paw luminance (b,c), standing Rt to Lt hind paw luminance (e,f), and distance traveled (g,h) (N = 4-6 mice; P values are determined by two-way ANOVA with Sidak correction in b,d and e; * p ≤ 0.05 for hG3BP1^K376Q^ vs. mCh, # p ≤ 0.05 for hG3BP1^K376Q^ vs. hG3BP1^K376R^, and ¶ p ≤ 0.05 for hG3BP1^K376R^ vs. mCh).

**Supplementary Figure 4:**
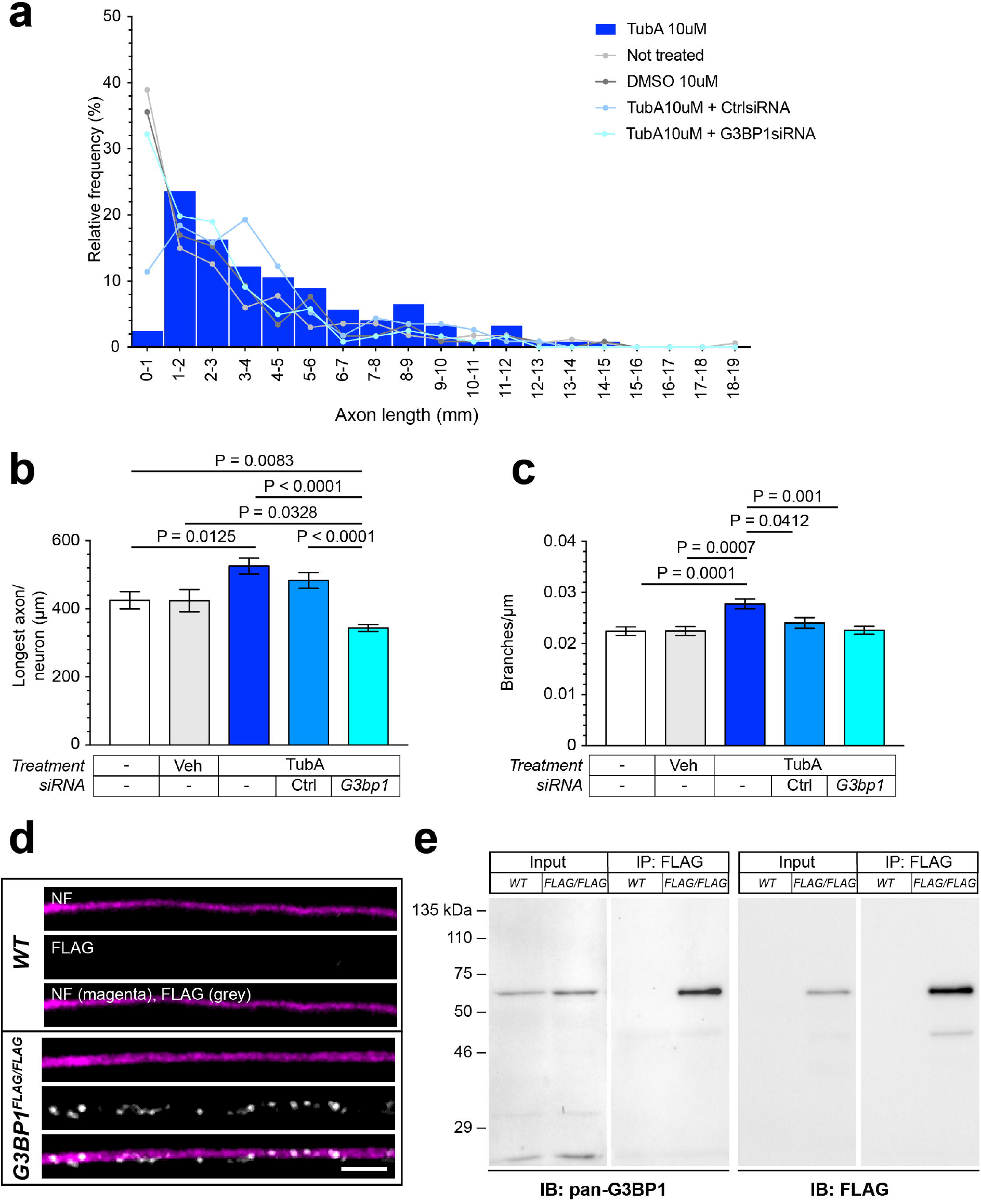
HDAC6 inhibitor Tubastatin A increases axon growth. **a)**, Histogram comparing axon length distributions in DRG neurons treated with vehicle or 10 µM TubA and siCntl or siG3bp1 from Figure 5b-c (1000 µm bins). **b-c)**, Longest axon per neuron (**b**) and axon branching (**c**) for DRGs transfected with vehicle or 10 µM TubA and siCntl or siG3bp1 from Figure 5b-c. **d)**, Representative confocal images for anti-NF + anti-FLAG immunostaining for DRGs cultured from WT and *G3BP1*^*FLAG/FLAG*^ mice [Scale bar = 10 µm]. **e)**, Immunoblots for anti-G3BP1 vs. anti-FLAG for protein lysates and anti-FLAG immunoprecipitates from WT and *G3BP1*^*FLAG/FLAG*^ mice DRG cultures (N ≥ 115 neurons across 3 independent cultures in a-c; P values by one-way ANOVA with Tukey’s post-hoc test in b-c).

**Supplementary Figure 5:**
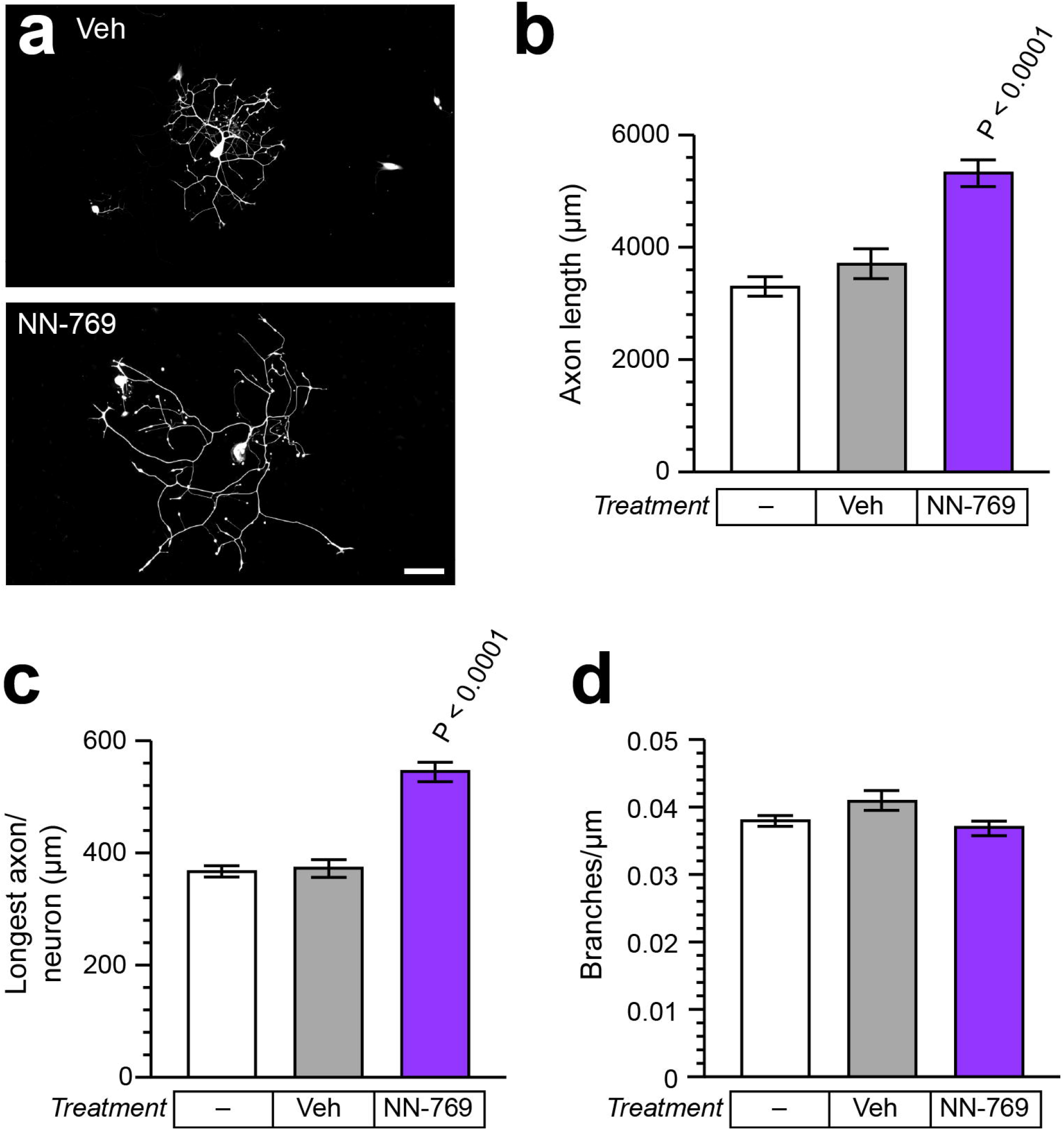
HDAC inhibition increases axon growth. **a)**, Representative images of DRG neurons cultured with HDAC6 inhibitor NN-769 vs. vehicle control (Veh; DMSO). **b)**, Quantification of total axon length of DRGs treated with 10 nM NN-769 for 2 h vs. Veh. **c-d)**, Longest axon length per neuron (**c**) and axon branching (**d**) for DRG cultrues treated as in a (N ≥ 180 neurons per condition across 3 replicated cultures in b-d; P values by one-way ANOVA with Tukey’s post-hoc in b-d).

**Supplementary Figure 6:**
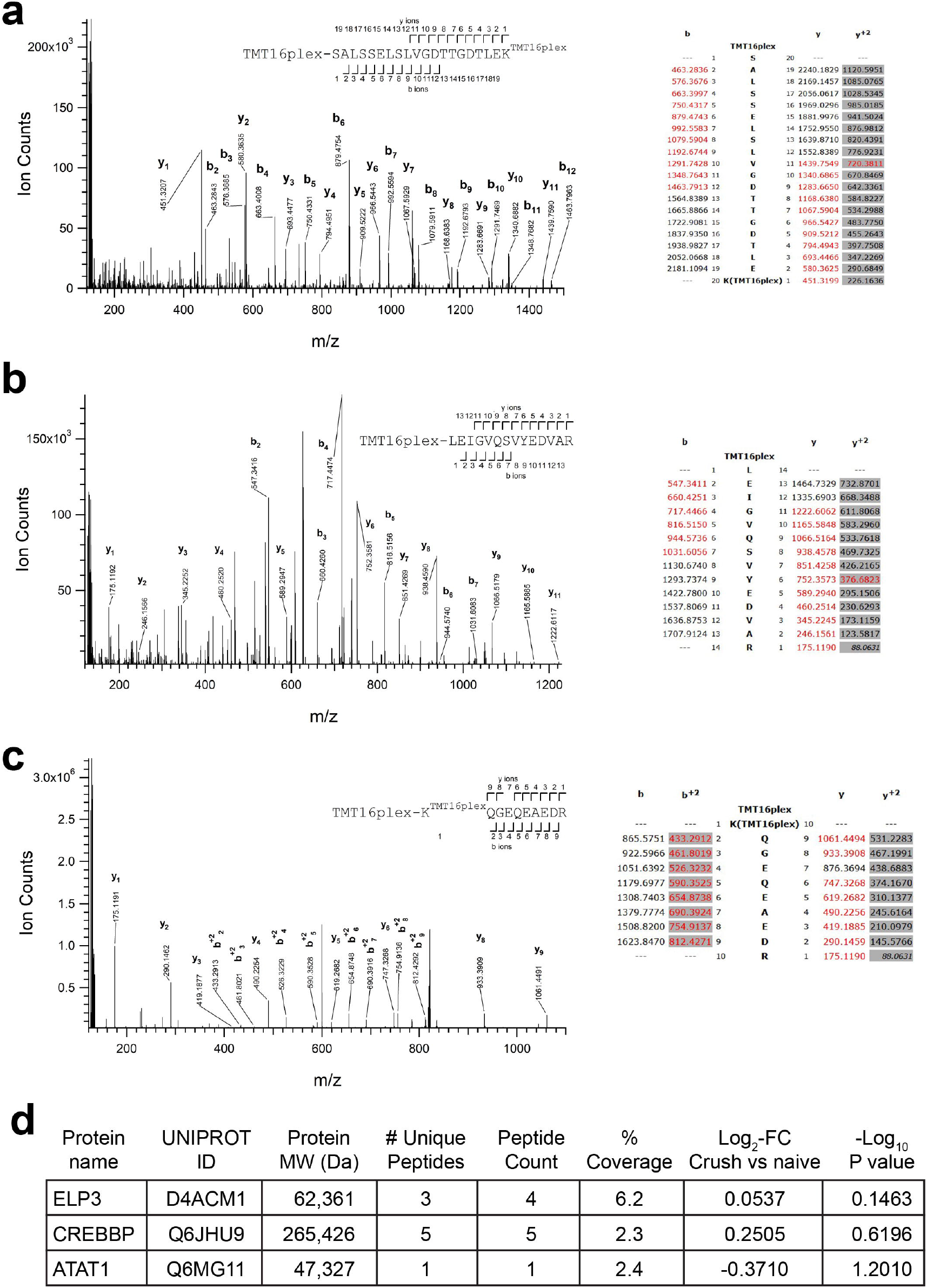
Mass spectrometry Identification of lysine acetyl transferases in rat axoplasm tryptic digests labelled with tandem mass tags reagents (TMT). **a-c)**, MS/MS spectra belonging to tryptic peptides spanning amino acids S2418 to K2437 of CREBBP (**a**), L243 to R256 of ELP3 (**b**), and K355 to R364 of ATAT1 (**c**), obtained by HCD fragmentation of precursor ions 878.1461^+3^, 628.0127^+3^, and 599.9908^+3^, respectively, of sciatic nerve axoplasm. Experimental masses of sequence ion peaks are labelled in the spectra, also indicating the fragment type (Roepstorff-Fohlmann-Biemann ions nomenclature). The position of the fragmentation events generating these ions are indicated in the sequences over the spectra. A table indicating the theoretical masses of the sequence ions according to the proposed modified sequences is shown on the right side of panels a-c, with red font indicating observed fragments. K (^TMT16plex^), TMT ε-amino modified Lysine; amino termini of these peptides is TMT modified (labelled in the figures as TMT16^plex^-). **d)**, Summary of LC-MS/MS data showing protein ID, peptide hits, and fold change for 7 d crush vs. naïve sciative nerve axoplasm.

**Supplementary Figure 7:**
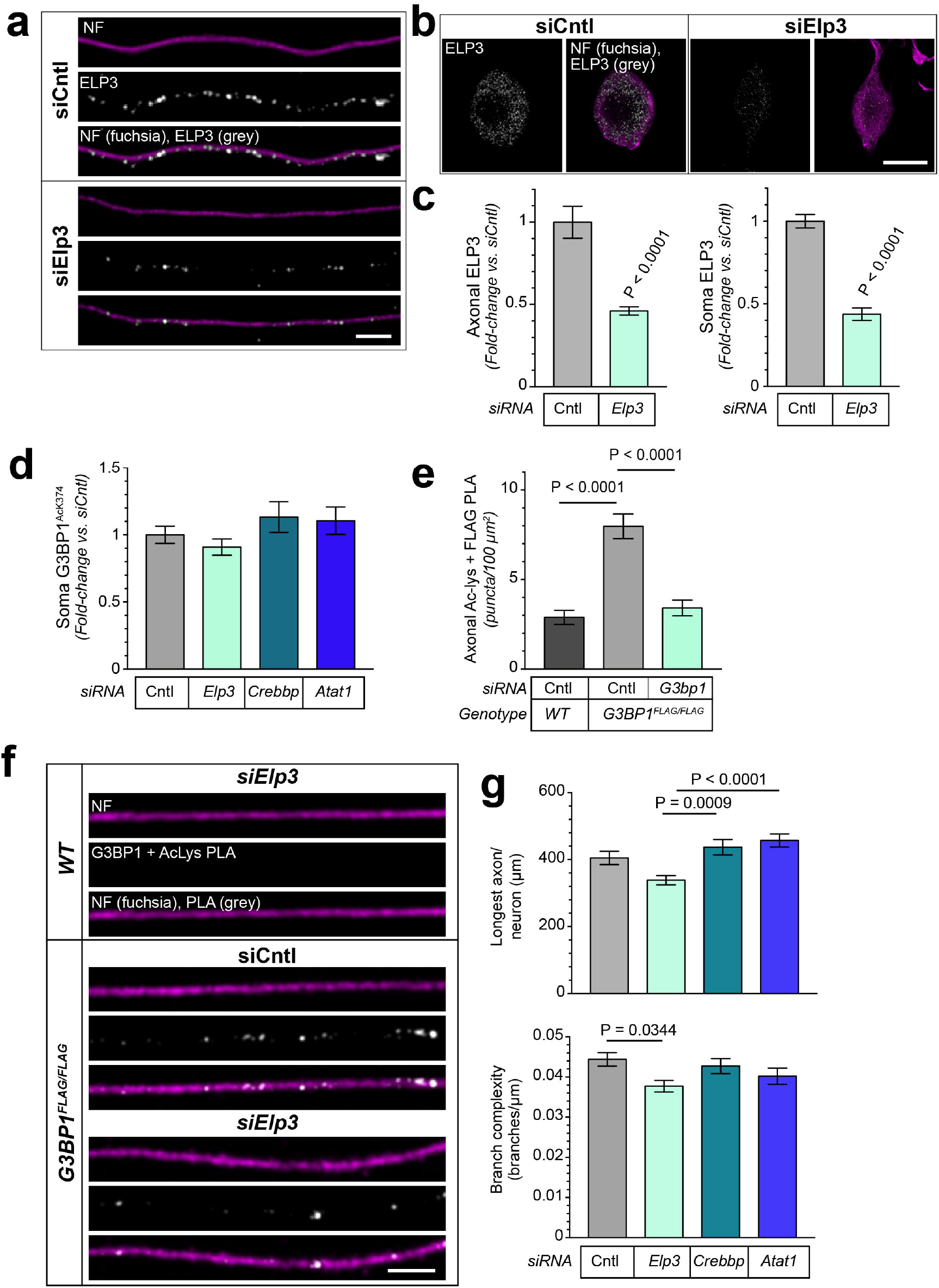
ELP3 depletion alters axon growth. **a-c)**, Representative confocal images for anti-NF + -ELP3 immunostaining in axons (**a**) and soma (**b**) for siCntl vs. siElp3 transfected DRG cultures. **c** shows quantitation of axonal and soma ELP3 levels across multiple cultures [Scale bars = 10 µm in a and 25 µm in b]. **d)**, Soma G3BP1^AcK374^ levels in siCntl, siElp3, siCrebbp, and siAtat transfected DRG cultures as in Figure 6b-c. **e-f)**, Mean PLA puncta density ± SEM in axons for anti-Ac-Lys + -FLAG in *WT* vs. *G3BP1*^*FLAG/FLAG*^ DRG cultures transfected with siCntl vs. siElp3 as indicated (**e**). Panel **f** shows representative confocal images of DRG cultures from e [Scale bar = 10 µm]. **g)** Longest axon per neuron (top) and axon branching (bottom) for siCntl, siElp3, siCrebbp, and siAtat transfected DRG cultures from Figure 6e (N ≥ 144 cell bodies and ≥ 36 axons across 3 independent cultures in c, ≥ 44 in d, 172 in e, and 178 in g; P value determined by Student’s t-test in c and one-way ANOVA with Tukey’s post-hoc in d, e, and g).

**Supplementary Figure 8:**
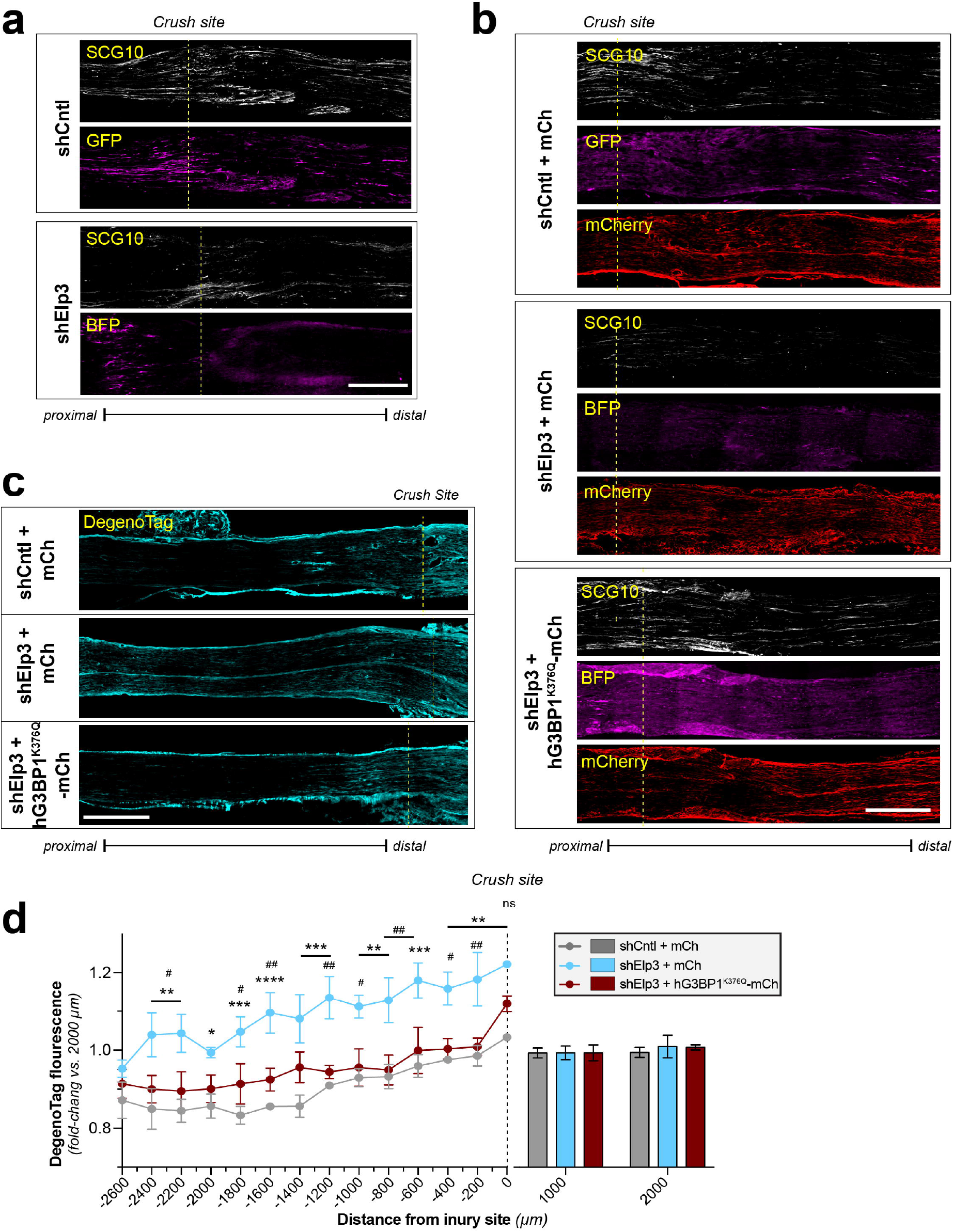
ELP3 depletion leaves axons susceptible to degeneration. **a)**, Representative SCG10 and BFP immunostaining of 5 d post-crush sciatic nerves from mice transduced with AAV-shCntl-GFP vs. -shElp3-BFP 14 d prior to nerve crush (19 d prior to analysis). Dashed yellow lines indicate injury site with proximal to the left [Scale bar = 500 µm]. **b)**, Representative SCG10, BFP and mCherry immunostaining of 5 d post-crush sciatic nerves from mice cotransduced with AAV-shCntl-GFP + -mCh vs. AAV-shElp3-BFP + -mCh or vs AAV-hG3BP1^K376Q^-mCh 14 d prior to nerve crush (19 d prior to analysis). Dashed yellow lines indicate injury site with proximal to the left [Scale bar = 500 µm]. **c)**, Representative *DegenoTag* immunostaining of 5 d post-crush sciatic nerves from mice transduced with AAV-shCntl-GFP vs. -shElp3-BFP 14 d prior to nerve crush (19 d prior to analysis). Dashed yellow lines indicate injury site with proximal to the left [Scale bars = 500 µm]. **d)**, Quantitation of *DegenoTag* fluorescence intensity of sciatic nerve neurons cotransduced with AAV-shCntl-GFP + -mCh vs. AAV-shElp3-BFP + -mCh or hG3BP1^K376Q^-mCh is shown as mean degeneration index ± SEM for each time point (N = 5-6 mice for a and b and 3 for d; ** p < 0.01, *** P < 0.005, and *** p < 0.001 for shCntl + mCh vs. shElp3 + mCh and # P < 0.05 and ## P < 0.01 for shElp3 + mCh vs. shElp3 + hG3BP1^K376Q^-mCh by two-way ANOVA followed by Sidak correction.

### SUPPLEMENTAL VIDEOS

**Supplementary Video 1: *Restoration of Digit Extension by G3BP1 acetylmimetic after sciatic nerve crush***. Representative video sequence acquired using the BlackBox One instrument showing mice transduced with AAV-hG3BP1^K376R^-mCh (left) and AAV-hG3BP1^K376Q^-mCh (right) at 10 d post-sciatic nerve crush. Differences in digit extension recovery are observed between the two conditions.

**Supplementary Video 2: *ELP3-depleted proximal axons rapidly degenerate after transection***. Time-lapse video of transected DRGs spot cultures transduced with AAV-shCntl-GFP (top) vs. AAV-shElp3-BFP (bottom). Images were acquired over 16 h at 10 min intervals and displayed at 7 frames per second [Scale bar = 100 µm].

**Supplementary Video 3: *Acetylmimetic G3BP1 prevents degeneration of proximal axon in ELP3-depleted neurons after transection***. Time-lapse video of transected DRGs spot cultures cotransduced AAV-shCntl-GFP + AAV-mCh (top panel), AAV-shElp3-BFP + AAV-mCh (middle panel), or AAV-shElp3-BFP + -AAV-hG3BP1^K376Q^-mCh (bottom panel). Images were acquired over 16 h at 10 min intervals and displayed at 7 frames per second [Scale bar = 100 µm].

## Notes

### Competing Interest Statement

JLT & PKS are co-founders of Rinnerva Therapeutics, which focuses on G3BP1 therapeutics.
NN, PM, and PTG are cofounders of HDAX Therapeutics.

